# Activation-independent capture of free fatty acids at the bacterial cell envelope by BrtB

**DOI:** 10.64898/2026.03.17.712404

**Authors:** Amaranta Kahn, Cristina I.F. Sousa, João P. A. Reis, David A. Russo, Adriana Rego, Ricardo J.C.V Queirós, Marine Cuau, Julie A. Z. Zedler, Sandra A.C. Figueiredo, Paulo Oliveira, Pedro N. Leão

## Abstract

Fatty acids (FAs) are key metabolites in living organisms, shaping membrane architecture and helping cellular acclimatization to changing conditions. Bacteria synthesize FA *de novo* but can also reclaim them from membrane lipids or uptake them from the environment, usually converting free fatty acids (FFAs) into activated FAs that can then be further metabolized. In cyanobacteria, it is the acyl-acyl carrier protein synthetase (Aas) that activates exogenous FFAs. Yet, the cyanobacterial enzyme BrtB was recently shown to esterify, *in vitro*, FFAs directly onto abundant chlorinated glycolipids (bartolosides), generating bartoloside fatty acid esters (B-FAs). Whether this chemistry operates *in vivo*, where it occurs, and what its implications are for cell physiology has remained unclear. Here we show that in the cyanobacterium *Synechocystis salina* LEGE 06099, BrtB captures FFAs at the cell envelope without prior activation to generate B-FAs. We found that supplemented FFAs were converted into B-FAs within minutes. Strikingly, this response occurred with minimal changes in gene expression and little alteration of the extracellular proteome, consistent with a pathway already in place. Additionally, we observed that B-FAs can further be hydrolyzed into hydroxybartolosides – the levels of the latter metabolites increase in response to FA supply, suggesting a transient sequestration of FFA. Our results demonstrate that activation of FFA is not the only route to their cellular incorporation and identify a specialized-metabolite pathway that captures exogenous FFAs at the cyanobacterial cell envelope.

## Introduction

Fatty acids (FAs) are central metabolites in cellular structure and remodeling, and energy production. They are crucial for carbon and energy storage, often found in the form of neutral lipids or lipid droplets ^1,2^, and play roles in regulating protein trafficking and signaling ^3,4^. Yet, most FAs are primarily used in the synthesis of complex lipids, which are essential components of cell membranes ^5^. While every organism synthesizes FAs *de novo* and esterifies them into lipids, many can also directly incorporate or recycle them into new lipid species, saving energy and increasing plasticity, representing fundamental processes across the various domains of life ^6,7,8^. All characterized FA incorporation and recycling mechanisms described thus far involve prior activation of free FAs (FFAs) as acyl adenylates, acyl phosphates or thioesters ^8,9^. After activation, and depending on the organism and FFA chain length, FAs can enter β-oxidation to generate energy. Alternatively, they can be routed into the FA synthesis II (FASII) pathway for elongation and/or incorporation into lipids ^8^. In this context, the phylum Cyanobacteria offers unique insights into how cells build, modify, and manage FAs, as they present a number of remarkable features. One of them is their capacity to perform plant-like oxygenic photosynthesis, which requires considerable overall lipid, acyl chain composition and unsaturation remodeling in response to environmental cues. In diderm bacteria, the key enzyme for activation and incorporation of FFAs is the acyl-acyl carrier protein (acyl-ACP) synthetase (Aas) ^9,10^. In cyanobacteria, Aas has been characterized in *Synechocystis* sp. PCC 6803 (henceforth S6803) and *Synechococcus elongatus* PCC 7942 as the enzyme that converts FFA to acyl-ACP through direct activation (Figure 1A). Aas has also been implicated into mediating promiscuous transfer of FFAs across biological membranes ^11–13^. The cyanobacterial Aas enzymes characterized to date display broad substrate tolerance toward exogenous FFAs ^12^. Additionally, although most living organisms possess a β-oxidation pathway that degrades FFAs into acetyl-CoA, most cyanobacteria appear to lack it, making their FA metabolism distinct from that of many other bacteria ^11,14,15^. Hence, Kaczmarzyk and Fulda ^12^ hint that FFAs released during lipid turnover are recycled through Aas, enabling remodeling of membrane lipids in response to environmental change. Consequently, the incorporation of FAs into primary metabolism lipids or into secondary metabolites has been considered as the only plausible destination for imported and recycled FFAs in cyanobacteria ^15^.

**Figure 1.**
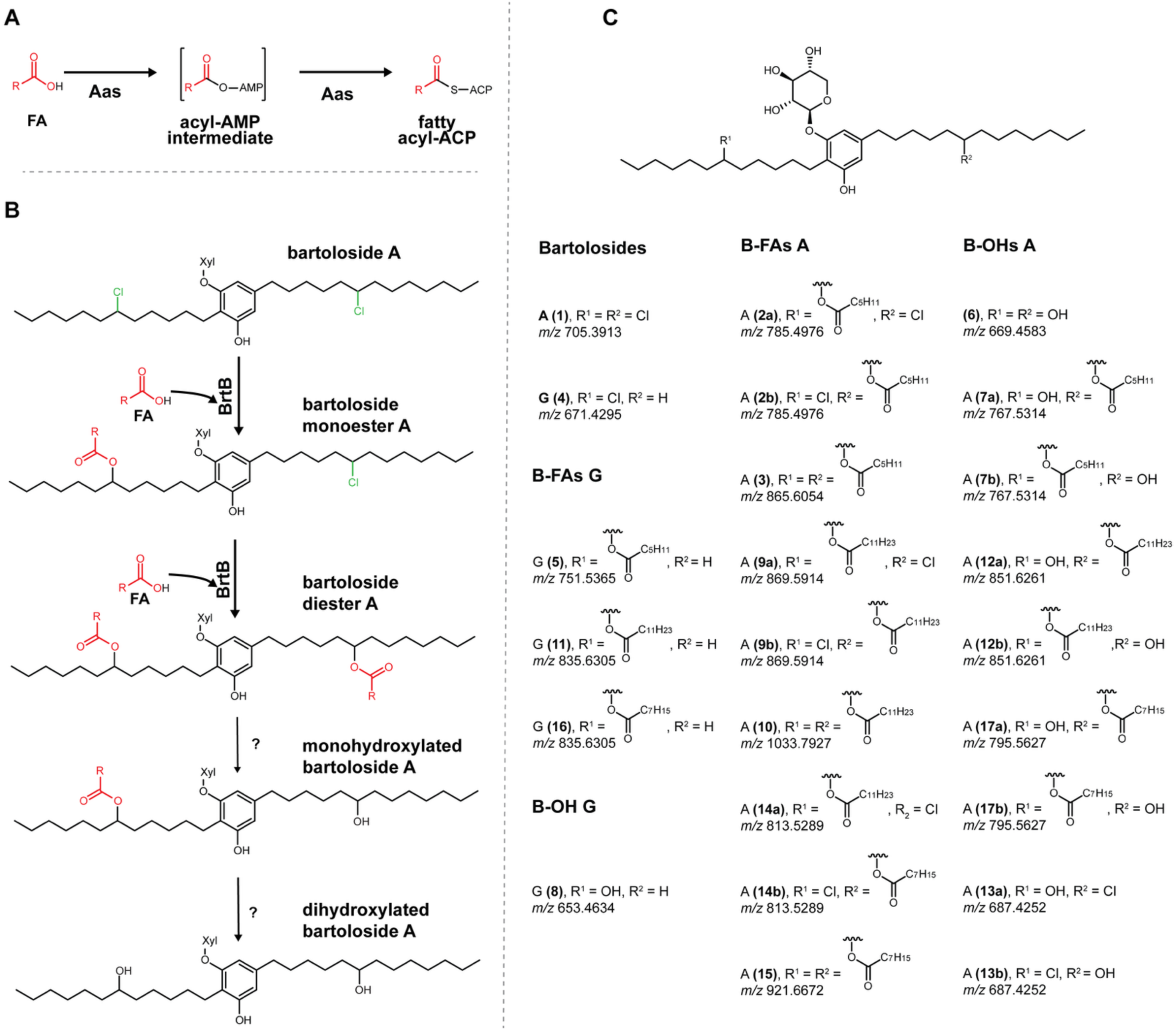
Free fatty acid (FFA) activation mechanism by acyl-acyl carrier protein (acyl-ACP) synthetase (Aas) and incorporation by BrtB, and bartoloside-related metabolites detected in this study. **(A)** FFAs incorporation by Aas. Aas catalyzes the reaction between a FFA and the terminal thiol of an ACP, leading to its loading onto the ACP. **(B)** FFA incorporation by BrtB. This carboxylate alkylating enzyme catalyzes O–C bond formation between a FFA and a secondary alkyl halide in a bartoloside A molecule to create a bartoloside monoester A. A second similar reaction on the remaining chlorine yields the diester. Subsequent hydrolysis by a putative esterase/lipase is proposed to generate monohydroxylated and then dihydroxylated bartolosides. **(C)** Generic bartoloside structure with variable substituents R^₁^ and R^₂^. Detected species are indicated, including bartolosides A (**1**) and G (**4**), the two most abundant bartolosides in S06099 ^17^; bartoloside fatty acid esters (B-FAs) derived from A and G bearing C6, C8, or C12 acyl chains (2, 3, 5, 9–11, 14–16) at R^₁^ and/or R^₂^; and hydroxybartolosides (B-OHs) derived from bartolosides A and G (**6–8, 12, 13, 17**) carrying OH at R^₁^ and/or R^₂^. Bartoloside A and bartoloside G share the same monoglycosylated dialkylresorcinol scaffold but differ in halogenation: bartoloside A is dichlorinated, whereas bartoloside G is monochlorinated, with one chlorine replaced by hydrogen ^14,17^. Reported *m/z* values correspond to the [M–H+HCOOH]^⁻^ formate adduct. FA, fatty acid; AMP, adenosine monophosphate; Xyl, xylose.

The bartolosides ^14,16^, a unique family of glycolipids that can be as abundant as 1% of the bacterial cell dry weight have been identified in eleven *Synechocystis salina* strains isolated from the Portuguese coast, and in *Synechocystis* sp. PCC 7338 isolated from Bodega Head (California, USA) ^17^. In *S. salina* LEGE 06099 (henceforth S06099), one of the highest yielding bartoloside producers, bartolosides were reported to undergo esterification with FFAs to create bartoloside fatty acid esters (B-FAs) ^17,18^. The gene cluster responsible for bartoloside biosynthesis (*brt*) ^16^ encodes the enzyme BrtB, responsible for the esterification step ^18^. *In vitro*, BrtB does not require activated FAs, directly esterifying FFAs with bartolosides. In this reaction, the FFA carboxylate acts as a nucleophile and attacks the alkyl chloride moieties present in the bartolosides, generating the ester bond with the displacement of HCl ^18^ (Figure 1B). In addition, it has also been shown that BrtB can esterify bartolosides *in vitro* with FAs ranging from C2 to C16, creating a large diversity of B-FAs. Metabolomic analysis of S06099 cells showed that a similar range of B-FAs occurs naturally, the most abundant of which are esters of bartolosides with saturated and unsaturated C16 and C18 FAs. In S06099, supplementation of culture medium with FFAs led to a drastic reduction of cellular bartoloside levels (>99%) and a concomitant increase in B-FAs ^18^. More recently, hydroxybartolosides (B-OHs) – in which the chlorinated position is replaced by an alcohol – have been reported in S06099, and were suggested to arise from B-FA hydrolysis ^17^ (Figure 1B).

In this work, we aimed to study the physiological relevance of BrtB. Using a combination of FA supplementation, stable-isotope tracing, transcriptomics and proteomics, we show that BrtB converts supplemented, non-activated FFAs into B-FAs *in vivo*, which are subsequently hydrolyzed to B-OHs. BrtB esterification of supplemented FFAs occurs within 30 min and is associated with only mild transcriptional changes. Exoproteome analyses support that BrtB is secreted and likely retained at the cell envelope, in agreement with rapid FFA incorporation into bartolosides prior to activation. Together, these data support a model in which direct FFA incorporation occurs at the S06099 cell envelope via BrtB-dependent bartoloside esterification and downstream formation of B- OHs. An overview of the bartoloside-derived metabolites tracked throughout the study, with compound numbering and *m/z* values, is provided in Figure 1C.

## Results

### BrtB captures exogenous fatty acids *in vivo* without prior activation

As BrtB has been demonstrated to generate bartoloside B-FAs *in vitro* without prior FA activation ^18^, we hypothesized whether BrtB would bypass FA activation by Aas in S06099 and directly incorporate them onto bartolosides ^8,12^. To test this, we followed the incorporation of ^18^O_2_-labeled FFAs by S06099 (Figure 2A; Figure supplement 1, Figure supplement 2A, Figure supplement 3A). We considered that if BrtB acts before the exogenously provided FFAs are activated by Aas, then we would observe two ^18^O atoms per esterification in B-FAs. Thus, we synthesized ^18^O_2_-labeled short-chain hexanoic acid, and medium chain dodecanoic acid (Figure 2B, Figure supplement 1, Figure Supplement 2B), which were used for the supplementation of S06099 cultures (Figure 2C, Figure supplement 2C, Figure supplement 3B). LC-HRESIMS analysis of the resulting cell extracts revealed the expected depletion of bartolosides ^17^, as well as mass features corresponding to ^18^O-labeled B-FAs (^18^O_2_-**2**, ^18^O_4_-**3**, ^18^O_2_-**5**, ^18^O_2_-**9**, ^18^O_4_-**10**, ^18^O_2_-**11**). Importantly, mass features corresponding to a single ^18^O atom incorporation per ester bond were not detected. The consistent incorporation of two ^18^O atoms per ester bond therefore indicates that the FFAs have not been previously activated, in agreement with the *in vitro* activity of BrtB ^18^ (Figure 2D, Figure supplement 2D, Figure supplement 3C-D).

**Figure 2.**
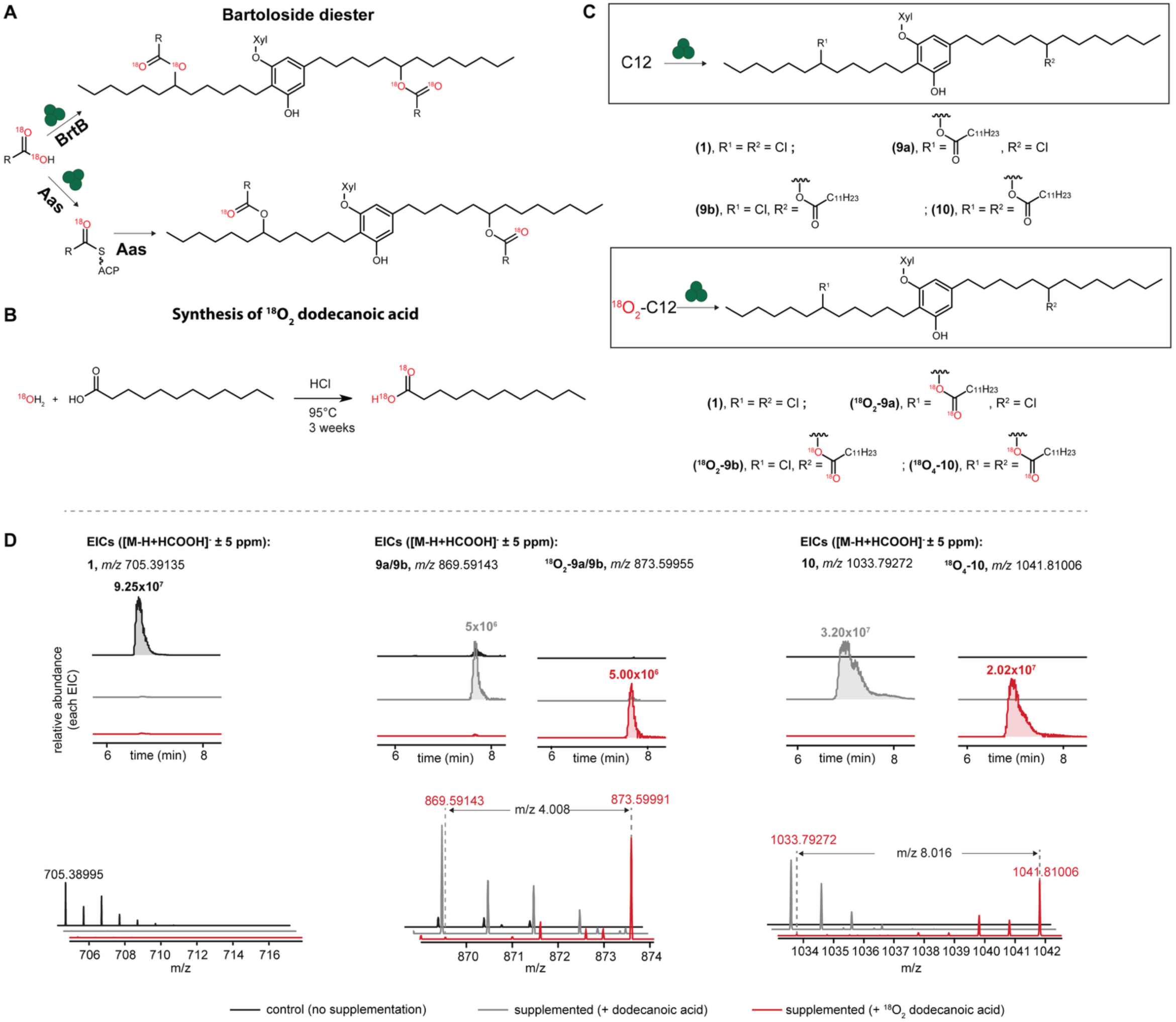
Supplementation of *Synechocystis salina* LEGE 06099 (S06099) with ^18^O_2_-labeled hexanoic acid (C6) leads to the direct esterification of bartolosides into bartoloside fatty acid esters (B-FAs). **(A)** Representation of the two possible pathways taken by the supplemented ^18^O_2_-C6 fatty acid (FA): direct incorporation by BrtB would lead to the incorporation of two (monoester) or four (diester) ^18^O atoms, while prior activation of the FA by the acyl-acyl carrier protein synthetase (Aas) would lead to the incorporation of one (monoester) or two (diester) ^18^O atoms. **(B)** Schematic representation of the synthesis of ^18^O_2_-hexanoic acid (C6) from H_2_^18^O and C6. **(C)** Structures of bartoloside A (**1**), mono- and dihexanoates esters (**2a**/**2b**, **3**), and their ^18^O- isotopologues generated upon supplementation of S06099 with ^18^O_2_-C6. **(D)** LC-HRESIMS-based detection of **1**, **2**, **3** and ^18^O_2 or 4_-labeled **2** and **3** in ^18^O_2_-C6 supplemented cultures of S06099. Values next to each peak in extracted-ion chromatograms (EICs) correspond to peak height (ion counts). ACP, acyl carrier protein; Xyl, xylose.

### B-FA levels plateau following FFA supplementation

Having established that B-FAs are formed without prior FFA activation, we next questioned whether BrtB could be saturated, and a plateau of B-FAs would be reached. To test this, we supplemented S06099 with three different concentrations (0.01 mM, 0.05 mM, 0.5 mM) of *d*_23_-dodecanoic acid and of *d*_15_-octanoic acid (Figure 3 and Figure supplement 4). Bartoloside-related species were then quantified 24 h and 7 days following supplementation (Source data 1). The LC-HRESIMS data were first screened for the relative abundance of bartolosides (**1**, **4**) and labeled B-FAs (*d*_23_-**9**, *d*_23_-**10**, *d*_46_-**10,** *d*_23_-**11**; *d*_15_-**14**, *d*_15_-**15**, *d*_30_-**15**, *d*_15_-**16**). As expected, 24 h post-supplementation, the level of bartolosides decreased proportionally to the concentration of supplemented FAs (Figure 3B, Figure supplement 4B). Surprisingly, the relative abundance of B-FAs increased to the same level regardless of concentration of supplemented FFAs (Figure 3C, Figure supplement 4C). The overall abundance of B-FAs decreased by half after 7 days, consistent with a degradation or modification mechanism. As for bartoloside levels, they increased about 3-fold after 7 days, likely from *de novo* bartoloside biosynthesis (Figure 3B-C, Figure supplement 4B-C).

**Figure 3.**
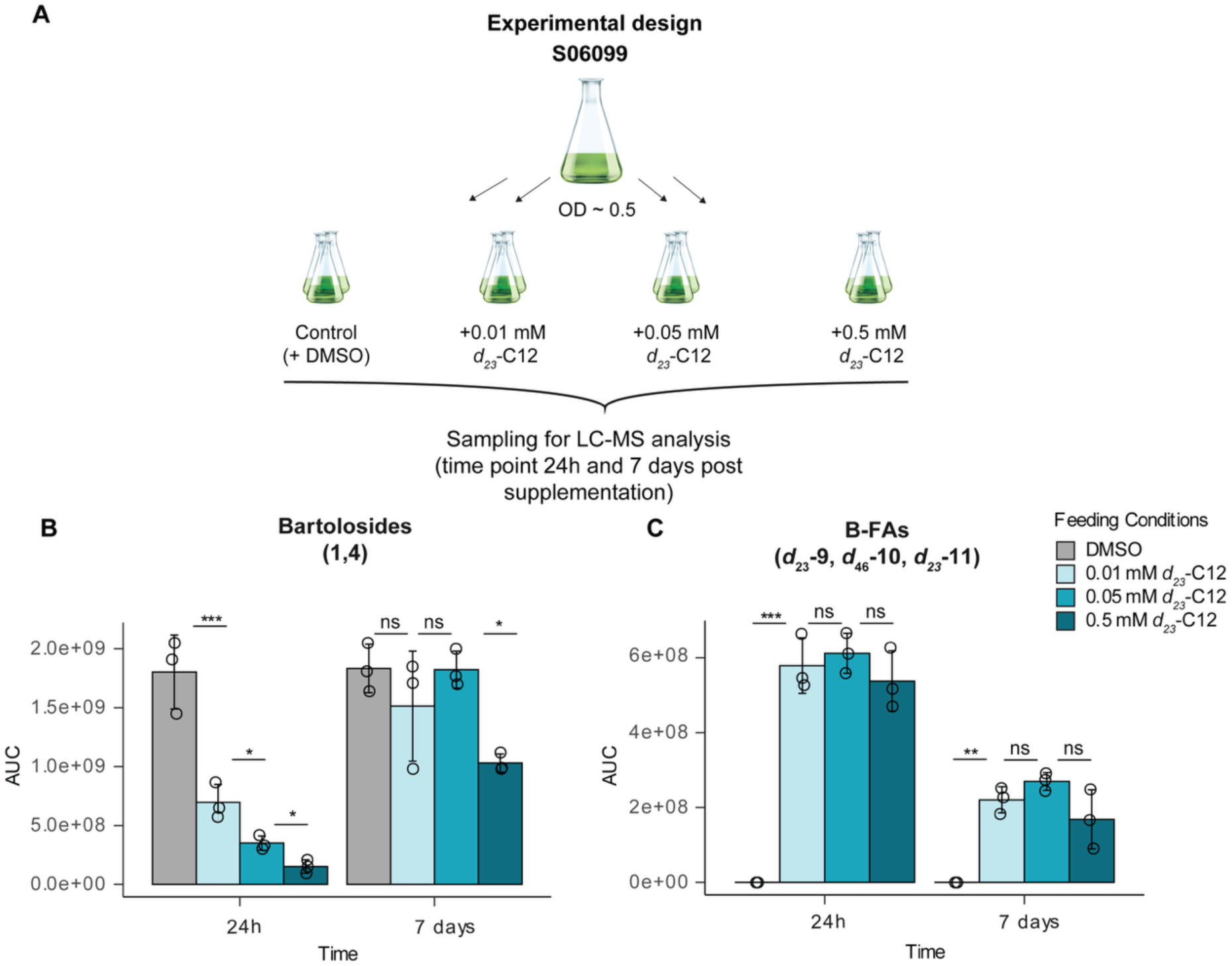
*Synechocystis salina* LEGE 06099 (S06099) bartoloside levels decrease with increasing supplementation of *d*₂₃–labeled dodecanoic acid (C12) while bartoloside fatty acid esters (B-FA) levels plateau. **(A)** Design of labeled fatty acid (FA) dose-response experiment. Mid-exponential cultures (OD ∼0.5) of S06099 were supplemented with 0.01, 0.05 or 0.5 mM *d*₂₃–C12 (or DMSO as a control) and harvested after 24 h and 7 days for LC-HRESIMS analysis. **(B)** Sum of area under the curves (AUCs) for **1** and **4**. **(C)** Sum of AUCs for *d*₂₃-labeled dodecanoate esters (B-FAs; *d*₂₃-**9**, *d*_46_-**10**, *d*₂₃-**11**). Bars show mean ± s.d. (n = 3 biological replicates); asterisks indicate significant differences between conditions (ns, not significant; * *p* < 0.05; ** *p* < 0.01; *** *p* < 0.001). All AUC values correspond to extracted ion chromatograms of the [M–H+HCOOH]^⁻^ formate adduct of the indicated species. DMSO, dimethyl sulfoxide; LC-MS, liquid chromatography-mass spectrometry.

### B-FAs can be hydrolyzed into B-OHs

Reis et al. ^17^ detected B-OHs in S06099 cells and postulated that these species likely arise from B-FA ester hydrolysis. To test this possibility, we revisited our stable isotope labeled oxygen supplementation assay data (Figure 2 and Figure supplement 2 and 3). Consistent with this hypothesis, ^18^O-labeled B-OH species ^18^O_2_-**6**, ^18^O_3_-**7** and ^18^O-**8** were detected in cultures supplemented with ^18^O_2_ hexanoic acid, while ^18^O_3_-**12** labeled B-OH species were detected in cultures supplemented with medium chain ^18^O_2_ -**12**-dodecanoic acid (Figure 4, Figure supplement 5). Upon ^18^O_2_ labeled and non-labeled hexanoic acid supplementation, B-FA and B-OH species (labeled and non-labeled, respectively) were detected at comparable relative abundance. In contrast, following ^18^O_2_-labeled and non-labeled dodecanoic acid supplementation, B-FA levels exceeded those of B-OH by approximately 20 to 30-fold respectively. We did not detect labeled chlorinated B-OH species (^18^O_2_**-9**) in none of the supplementation experiments. In accordance with previous work^17^, we also detected non-labeled B-OHs (**6**, **12**) without FFA supplementation (Figure 4C, Figure supplement 5C). This seems to indicate that FAs are not exclusively recycled from B-FAs formed upon FA supplementation, but also from B-FAs formed with FFAs resulting from cellular metabolism. Overall, our *in vivo* data suggest that exogenously provided FFAs are esterified with bartolosides prior to activation, and that the formed B-FAs can be hydrolyzed into B-OHs.

**Figure 4.**
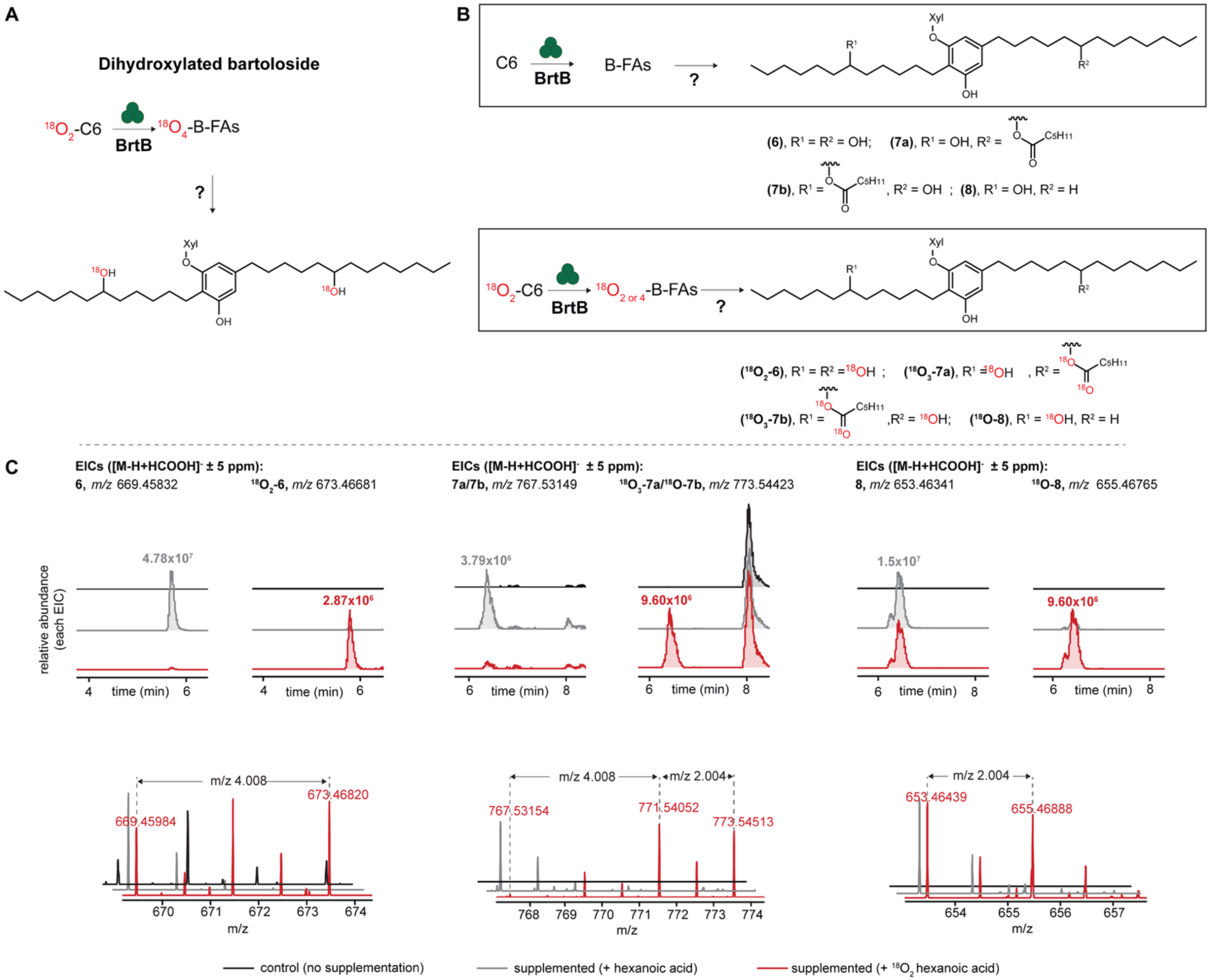
Bartoloside fatty acid esters (B-FAs) are hydrolyzed to hydroxybartolosides (B-OHs). **(A)** Representation of proposed events following ^18^O_2_-hexanoic acid (C6) supplementation: direct esterification by BrtB leading to the incorporation of two or four ^18^O atoms (two per each ester bond), followed by subsequent hydrolysis of the ester bonds to generate dihydroxylated bartolosides containing one or two ^18^O atoms. **(B)** Structures of B-OHs (**6**, **7a**/**7b**, **8**) and their ^18^O-isotopologues generated upon supplementation of *Synechocystis salina* LEGE 06099 (S06099) with ^18^O_2_-C6. **(C)** LC-HRESIMS-based detection of **6**, **7**, **8** and labeled **6**, **7**, **8** in ^18^O_2_-C6 supplemented cultures of S06099. Values next to each peak in extracted ion chromatogram correspond to peak height (ion counts). Xyl, xylose.

### B-FAs undergo increased hydrolysis upon FA supplementation

To understand whether B-FA hydrolysis could explain the stable relative abundance of B-FAs independent of increasing concentration of supplemented FFAs (Figure 3C), we revisited the data from the stable isotope *d*_23_-dodecanoic acid and *d*_15_-octanoic supplementation experiment. We tracked the B-OH species that could be deuterated (*d*_23_-**12**; *d*_15_-**17**) as well as the non-labeled species **6**. As expected, we detected labeled and non-labeled B-OH species (**6**, *d*_23_-**12**; *d*_15_-**17**, source data 1). Moreover, we observed that the levels of B-OH did increase proportionally with supplemented FFA concentration after 24 h (Figure 5A-B), which suggests that B-FA hydrolysis could account at least partly to the dose-independence of B-FA accumulation across supplementation concentrations (Figure 3C). Hydrolysis of B-FAs appeared to continue at high levels for the remaining of the *d*_23_-dodecanoic acid experiment at the highest concentration only (Figure 5A), likely reflecting turnover and reduced pool sizes. To question the fate of those hydrolyzed FA, we searched for the incorporation of labeled FFAs into two membrane lipid classes: phosphatidylglycerols (PGs) and sulfoquinovosyl diacylglycerols (SQDGs). Indeed, we observed a consistent increase of labeled PGs and SQDGs with increasing concentration of *d*_23_-dodecanoic acid (Figure 5C-D, Source data 2). The exception was deuterated PGs levels 7 days post-supplementation, which showed high variability across replicates. *d*_23_-SQDGs levels decreased by half from 24 h to 7 days after supplementation. However, because high levels of labeled PGs and SQDGs were detected even for the lowest concentration of supplemented perdeuterated FFAs, it is not possible to ascertain the relative contributions to this glycolipid pool from FFAs activated through Aas that entered primary metabolism versus those originating from B-FA hydrolysis. The level of non-labeled glycolipids did not seem to vary with labeled-FA supplementation (Figure supplement 6), indicating that their synthesis was not impacted overall. To summarize, these results indicate that bartoloside esterification and hydrolysis establish a steady B-FA level. The FFAs released during B-OH formation likely contribute to the cellular FFA pool and could be reincorporated into membrane lipids, but further experiments are needed to confirm this.

**Figure 5.**
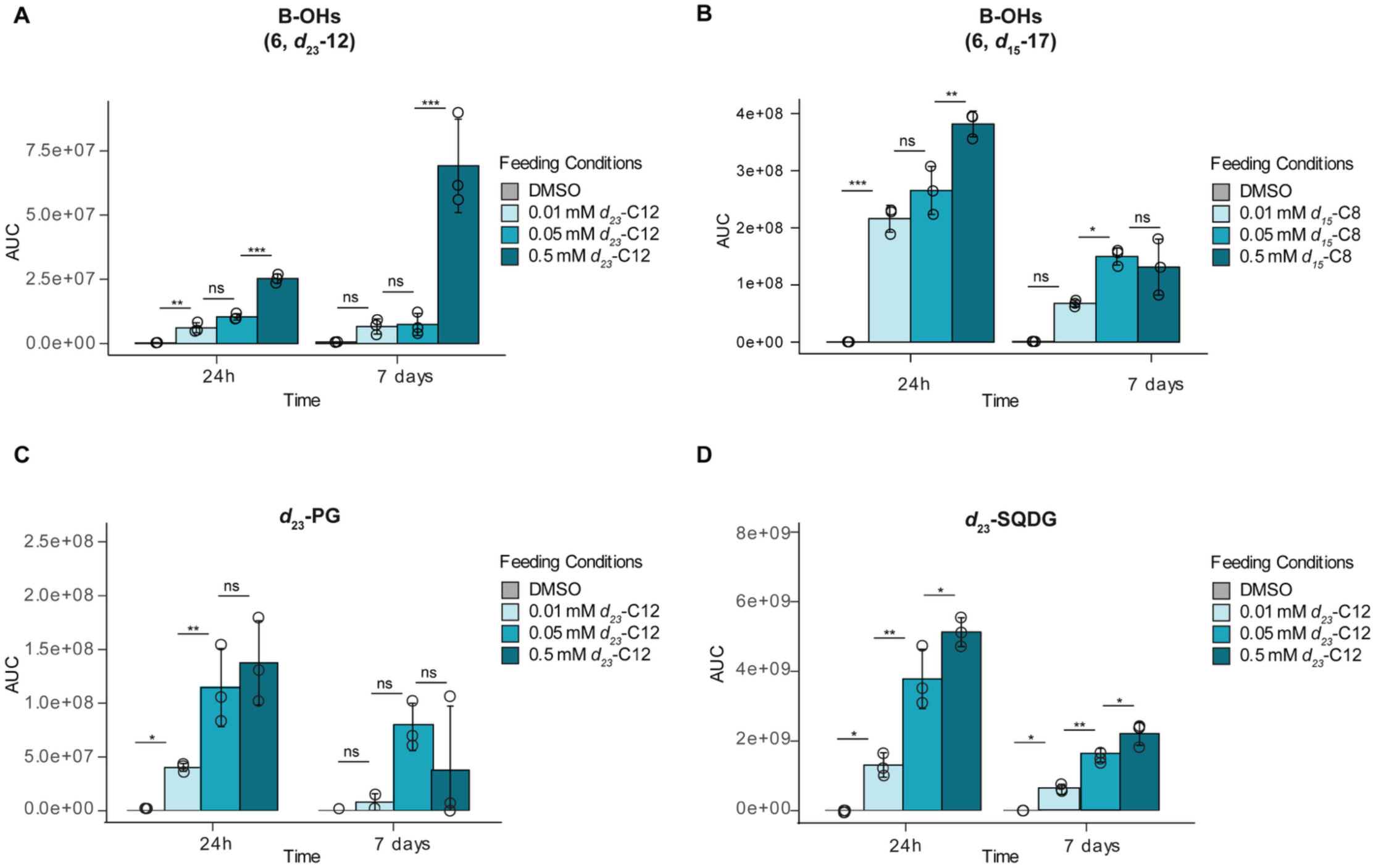
Hydroxybartoloside (B-OH) and anionic lipid levels vary with increasing supplementation of *d*₂₃–dodecanoic (C12) and *d_15_*–octanoic (C8) acids. Sum of area under the curves (AUCs) for: **(A)** non-labeled (**6**), *d*₂₃-labeled B-OHs (*d*₂₃-**12**) and **(B)** non-labeled (**6**) and *d*_15_-labeled B-OHs (*d*_15_-**17**). **(C–D)** Sum of AUCs for *d*₂₃-labeled phosphatidylglycerol (*d*₂₃-PG) and sulfoquinovosyldiacylglycerol (*d*₂₃-SQDG), respectively. Bars show mean ± s.d. (n = 3 biological replicates); asterisks indicate significant differences between conditions (ns, not significant; * *p* < 0.05; ** *p* < 0.01; *** *p* < 0.001). All AUC values correspond to extracted ion chromatograms of the [M–H+HCOOH]^⁻^ formate adduct of the indicated species. DMSO, dimethyl sulfoxide.

### B-FA acyl composition remains stable while glycerolipids remodel with temperature

An increase of labeled B-OH and anionic glycerolipids was observed with increasing FFA supplementation (Figure 5). Combined with the fact that basal levels of B-FAs are detected in non-supplemented cells^18^, this could indicate that the acyl chains from membrane lipids exchange with B-FA acyl chains. Temperature shifts are known to induce FA composition changes in membrane lipids ^19–21^. Having this in mind, we aimed to understand if such a response could be reflected in B-FAs acyl chain composition. To do so, we cultivated S06099 at 17 °C and at 30 °C and further collected samples at two time points (15 and 20 days). The chosen time points assure membrane composition stability and ensure one matching physiological state under the two different growth temperatures (OD ∼0.5). When grown at 17 °C and 30 °C, we observed at both time points that S06099 B-FA acyl composition remain unchanged (Figure supplement 7A, Source data 3). Whereas the anionic glycerolipids (PGs and SQDGs) acyl composition displayed the expected temperature-dependent remodeling ^19–21^ (Figure supplement 7B-C, Source data 3). More precisely, PGs showed a significant decrease in overall unsaturation at 30 °C relative to 17 °C and SQDGs displayed significant temperature-dependent decrease in both long-chain content and unsaturation. Thus, the major origin of B-FAs acyl chains does not seem to be membrane glycerolipids, and vice versa, at least under the tested conditions.

### FA supplementation does not induce major transcriptomic responses

Although membrane-lipid and B-FA acyl-chain compositions were not correlated, FFA supplementation caused B-FAs and B-OHs to accumulate in a dose-dependent manner. We therefore performed transcriptomics profiling after FFA supplementation to probe the cellular role of B-FAs and potentially identify key genes associated with the Brt system. Our previous experiments (Figures 2 to 5, Figure supplements 2 to 7) and previously published work ^18^ characterized bartoloside, B-FA, and B-OH formation in S06099 after extended incubations (24 h to a month). However, cyanobacterial transcriptional responses to environmental stimuli can begin within minutes and evolve over hours ^22–25^. Here, to assess how the S06099 transcriptome responds to the FFA supplementation stimulus, we initially searched for two time points that would capture the early rise in B-FA accumulation. We thus quantified cellular bartoloside-related metabolites after 30 min and 6 h of dodecanoic acid supplementation, to determine whether rapid and increasing bartoloside metabolic signatures occur in this cyanobacterium (Figure 6A, Source data 4). LC-HRESIMS analysis of cell extracts 30 min after dodecanoic acid supplementation revealed high levels of bartoloside-dodecanoic acid esters (B-FAs **9**, **10** and **11**), while the abundance of the bartolosides **1** and **4** were significantly lower than in the control. Six hours post-supplementation, bartoloside levels were roughly half of those in control cultures, whereas total B-FA levels approximately doubled compared with the 30-min time point (Figure 6B). The combined abundance of B-OH (**6**, **8**, **12**, **13**) was slightly lower relative to control cultures, but the dodecanoic acid-containing B-OHs (**12a/12b**) accumulated to high levels at both 30 min and 6 h only in supplemented cultures (Figure supplement 8).

**Figure 6.**
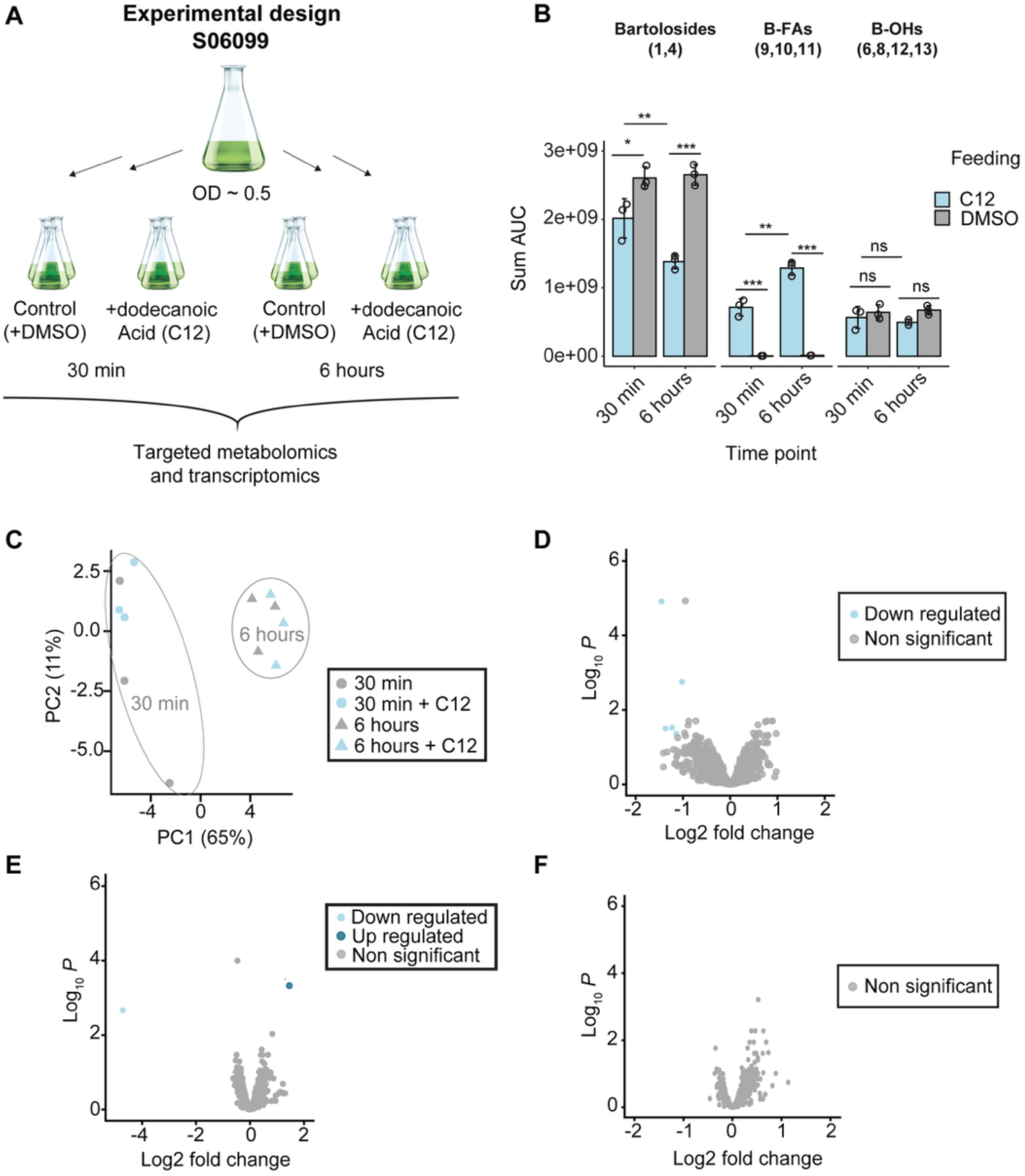
Dodecanoic acid (C12) supplementation remodels bartoloside-related species with minimal transcriptional response. **(A)** Experimental design. Mid-exponential *Synechocystis salina* LEGE 06099 (S06099) cultures (OD ∼0.5) were split into four conditions in biological triplicate (n = 3) and supplemented with a final concentration of 0.5 mM of C12 or DMSO alone (control); samples were collected at 30 min and 6 h for LC-MS profiling and RNA-seq. **(B)** Summed LC-MS peak area under the curve (AUC) for bartolosides (**1**, **4**), bartoloside fatty acid esters (B-FAs; **9**–**11**), and hydroxybartolosides (B-OHs; **6**, **8**, **12**, **13**) in dodecanoic-acid-treated versus control cultures at 30 min and 6 h; bars show mean ± s.d. (n = 3). Statistics: * *p* < 0.05, ** *p* < 0.01; ns, not significant. **(C)** Principal component analysis (PCA) of transcriptomes, colored by condition and shaped by time point. **(D–F)** Volcano plots log_2_ fold change versus –log_10_ adjusted *p* value (FDR) for dodecanoic acid versus DMSO at 30 min **(D)**, 6 h **(E)**, and both time points combined **(F)**; Genes significantly differentially expressed [when |log_2_ fold change| ≥ 1 and adjusted *p* value (FDR) ≤ 0.05] are highlighted. DMSO, dimethyl sulfoxide.

After establishing that significant bartoloside metabolic changes were observed within the defined timescale, we then investigated the effect of dodecanoic acid on the expression of S06099 genes by RNAseq transcriptomics analysis, 30 min and 6 h post-supplementation. Principal component analysis (PCA), based on the overall expression profile, showed a clear separation between the two-time points, as the samples clustered together independently of FA supplementation (Figure 6C). Pairwise Pearson correlations were >0.99 for all library-to-library comparisons, including across time points, indicating highly consistent transcriptome profiles across samples (Figure supplement 9). Regardless of supplementation, the 30 min and the 6 h samples showed consistently high pairwise correlations and clustered by time rather than treatment (Figure supplement 9). We nevertheless searched for differentially expressed (DE) genes, with a focus on FA metabolism and the *brt* gene cluster. Genes were considered significantly DE when |log2 fold change| ≥ 1 and adjusted p-value ≤ 0.05. Surprisingly, and despite the clear metabolic response in bartoloside levels, the RNAseq analysis identified only six and eight DE genes at 30 min and 6 h following supplementation, respectively (Figure 6D and E, Table 1), with no overlap between time points (Figure 6F). Sequence-based annotation and homology searches indicated that neither the *brt* genes ^18^ nor any obvious FA metabolism genes were DE upon supplementation (Table 1). None of the DE genes had assigned KEGG Orthology or Gene Ontology terms, preventing enrichment analysis. To test whether the transcriptional response depends on BrtB activity, we attempted to generate a Δ*brtB* mutant in S06099. Despite repeated natural transformation, electroporation, and conjugation-based approaches, we did not obtain stable mutants, preventing transcriptomic analyses in a BrtB-deficient background.

**Table 1.** Differentially expressed genes at 30 min and 6 h after dodecanoic acid (C12) supplementation.

| Time point | Gene ID<sup>a</sup> | LFC<sup>b</sup> | $p$-value | Annotation<sup>c</sup> |
| --- | --- | --- | --- | --- |
| **30 min** | 14_4 | -1.02 | 1.76E-03 | Glycosyltransferase |
|  | 14_114 | -1.14 | 4.38E-02 | FkbM family methyltransferase |
|  | 47_97 | -1.23 | 2.99E-02 | LabA-like NYN domain-containing protein |
|  | 50_81 | -1.37 | 3.14E-02 | Protein of unknown function (DUF433) |
|  | 16_22 | -1.45 | 1.23E-05 | Response regulator (cheY-homologous receiver domain) |
| **6 h** | 21_10 | 1.46 | 4.17E-04 | Heterocyst frequency control protein PatD-like |
|  | 43_1 | -4.71 | 2.15E-03 | Cell wall-associated hydrolase family protein (fragment) |

<sup>a</sup>Gene identifiers correspond to the ORF/locus IDs used for annotation of the *Synechocystis salina* LEGE 06099 genome and match the IDs provided to eggNOG-mapper. <sup>b</sup>Log<sub>2</sub> fold change (supplemented vs control). <sup>c</sup>Functional annotations were assigned with eggnog-mapper v2 and refined using InterProScan and BLASTp.

Across both time points, the DE genes (six at 30 min; eight at 6 h) are predicted to encode a glycosyltransferase, an FkbM-family methyltransferase, a CheY-like receiver-domain response regulator, a LabA-like NYN-domain protein, a DUF433 domain-containing protein, and a truncated cell-wall–associated hydrolase fragment (Table 1). One upregulated gene carried a PatD-like annotation by homology, which in a non-heterocystous strain likely reflects conserved domains rather than a heterocyst-specific role. In contrast, consistent with time of day or circadian regulation in cyanobacteria ^26,27^, 135 genes were DE between 30 min and 6 h irrespective of treatment (Figure supplement 10; Supplemental table 1). Overall, bartoloside-related metabolites changed as soon as 30 min after FFA supplementation, whereas the transcriptome showed minimal treatment-associated variation, consistent with a constitutive enzymatic response rather than transcriptional adaptation.

### BrtB is found in the extracellular medium and is likely associated to the cell envelope

The near absence of DE genes upon FFA supplementation contrasted with the strong bartoloside-associated metabolic response. Because BrtB-mediated incorporation also bypasses Aas-dependent activation, we considered that BrtB might physically access FFAs before Aas could. And that this could also be linked to a reduced early transcriptional response. Thus, we examined BrtB subcellular localization. In S6803, Aas has been detected in both total membrane and cytosolic fractions, although its distribution among individual membrane systems remains unclear ^28^. Inspection of the BrtB primary sequence revealed features consistent with type I secretion system substrates (Text supplement 1),^29^ including the absence of a canonical N-terminal signal peptide, predicted Ca²⁺-binding motifs, an unusually acidic predicted pI (3.73), and an amino acid composition rich in glycine (12%) and with no cysteines. To test whether BrtB is secreted across the cell envelope, we analyzed the S06099 exoproteome. Using the EXCRETE workflow^29^, BrtB was identified as the second most abundant protein among 94 proteins detected in the S06099 exoproteome (Figure 7A; Source data 5).

**Figure 7.**
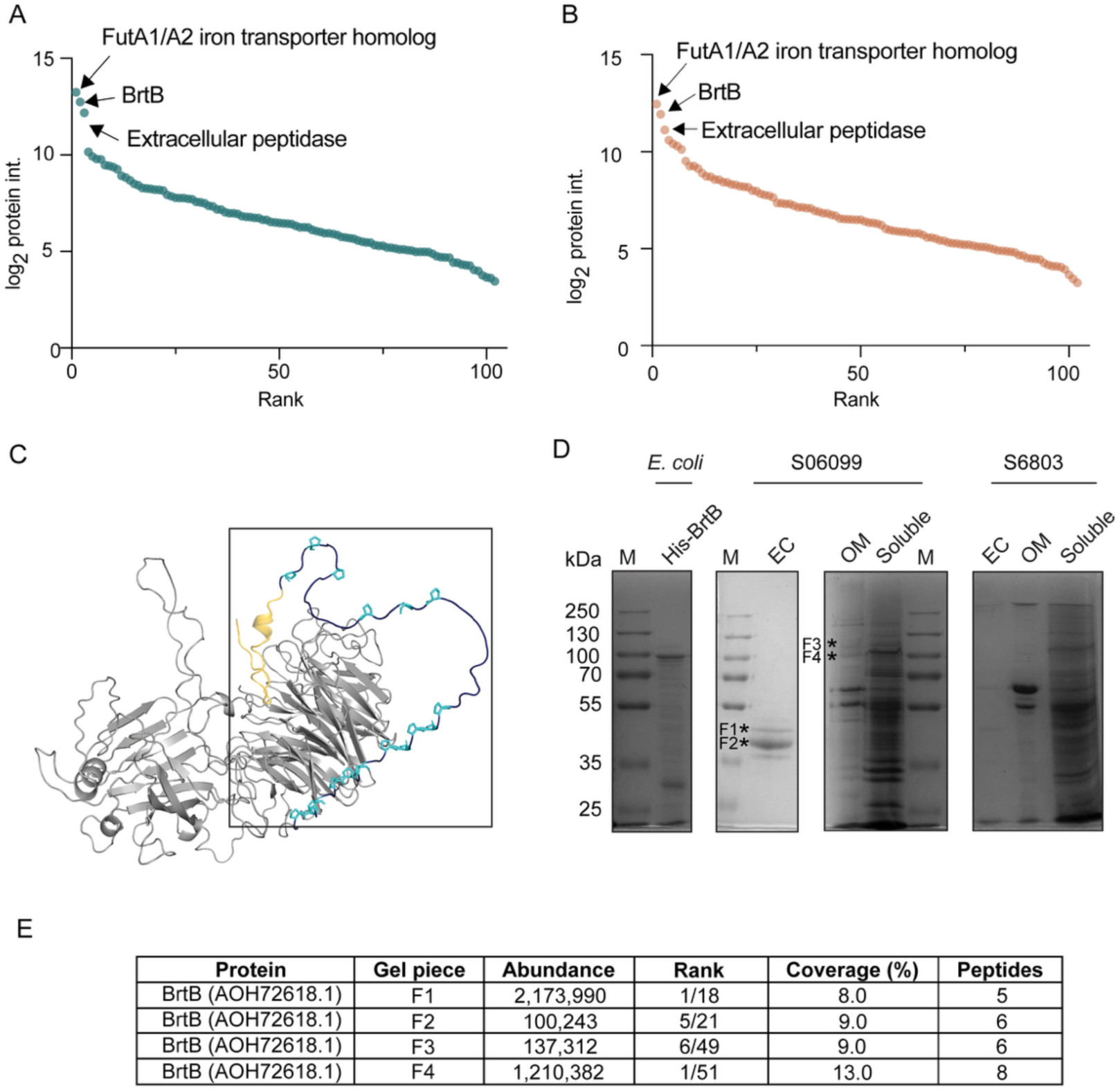
BrtB localizes in the cell envelope of Synechocystis salina LEGE 06099 (S06099). (A, **B)** Intensities of the proteins identified in the S06099 exoproteome in the control **(A)** and after hexadecanoic acid supplementation **(B)** ordered by rank. The three most abundant proteins are labeled. **(C)** Predicted structure of BrtB from the AlphaFold Protein Structure Database (AF-A0A1B3Z488-F1-v4), rendered in PyMOL. The β-propeller region (residues 1–752) is shown in grey; the C-terminal region (residues 753–831) is shown in dark blue, with prolines within this region highlighted in cyan (sticks) and the GS linker (residues 809–831) highlighted in yellow. The C-terminal region is additionally outlined with a black box. **(D)** Coomassie blue-stained SDS-polyacrylamide gels showing: purified BrtB expressed in *Escherichia coli* with a N-terminal histidine tag (His-BrtB; 1.5 µg of protein was loaded) – left-hand side panel; the cell free extracellular fractions (EC), outer membrane fractions (OM), and soluble fractions of S06099 (central panels), and of *Synechocystis* sp. PCC 6803 (S6803) (right-hand side panel). Cultures were normalized to the same cell density (OD_750_) so that comparable amounts of total protein could be loaded for S06099 and S6803. Bands excised for LC-MS/MS analysis are indicated by asterisks and labeled F1–F4. M, molecular mass marker; molecular masses (kDa) are indicated on the left. **(E)** Proteomic summary of BrtB (AOH72618.1) identified from gel pieces **F1–F4**, reported normalized abundance, rank among non-contaminant proteins, sequence coverage (%), and number of identified peptides. The full quantitative dataset is provided in Data Source 5-8, and the ten most abundant non-contaminant proteins identified in each gel piece are listed in Table Supplement 2.

We next asked whether FFA supplementation alters the abundance of secreted proteins, including BrtB. To reflect the physiology of S06099, we supplemented cultures with hexadecanoic acid (C16), a major fatty acid in S6803 membrane lipids ^30^ and a prevalent acyl chain in S06099 B-FAs ^18^. After 24 h supplementation, we did not detect changes in secreted protein abundance, and BrtB remained the second most abundant protein in the exoproteome (Figure 7B, Source data 5).

Given the abundance of BrtB in the exoproteome, and the fact that it contains a predicted outer-membrane (OM) anchor domain (Figure 7C), we assessed BrtB localization to the OM. We prepared cytoplasmic, OM, and cell-free extracellular protein fractions from both S06099 and S6803 and analyzed them by SDS-polyacrylamide gel electrophoresis. S6803 lacks the *brt* gene cluster and served as a control. In the OM fraction of S06099 we observed bands at an apparent molecular weight compatible with full-length BrtB (≈ 90 kDa) (Figure 7D, F3 and F4). We then excised the corresponding OM bands, as well as prominent bands present in the extracellular fraction of S06099 but not S6803, and analyzed them by LC–MS/MS. BrtB peptides were detected in both the OM and extracellular fractions (Figure 7E; Source data 6–9, Table supplement 2), with BrtB being the most abundant protein identified in bands F1 and F4; the extracellular BrtB peptides were detected in bands migrating at ≈40–50 kDa, consistent with BrtB-derived fragments, likely arising from BrtB proteolysis. Additional proteins detected in the OM-associated bands included canonical OM components, such as BamA/TamA family OM proteins and TonB-dependent receptor proteins, as well as other predicted β-barrel proteins (Table supplement 2).

Overall, BrtB is among the most abundant proteins detected in the S06099 exoproteome and is also detected in the OM fraction. Its abundance was not altered by FFA supplementation, consistent with an enzyme that is constitutively present at the cell envelope rather than upregulated under these conditions.

We next examined whether S06099 displays envelope ultrastructural features consistent with an expanded outer-envelope compartment, which could provide a physical context for BrtB localization. Transmission electron micrographs showed that S06099 shares major ultrastructural characteristics with the phylogenetically close strain S6803, including similar cell width in cross-section (∼1.7 µm diameter in both strains), thylakoid organization, cytoplasmic inclusions (e.g., glycogen granules), and an apparent S-layer (Figure supplement 11). However, in multiple independent preparations and across multiple non-overlapping fields of view across each grid, we observed several reproducible differences in the envelope region. In particular, in TEM thin sections, we measured a larger apparent OM-to–S-layer spacing in S06099 (78 ± 13 nm) than in S6803 (48 ± 7 nm) and S06099 displayed vesicle-like structures in the space between the OM and the S-layer as well as occasional surface protrusions (Figure supplement 11). These differences were observed irrespective of hexadecenoic acid supplementation, and an eventual link to the bartolosides or BrtB activity cannot be established or ruled out without follow-up experiments.

## Discussion

Uptake and recycling of FFA in bacteria have so far been described to occur only after activation as acyl-ACP, acyl-CoA, or acyl-phosphate ^8,11,12^. Outside bacteria, CoA-independent transacylases and cholesterol esterifying enzymes ^33,34^ support that acyl transfer, including esterification, can proceed without high-energy thioesters ^31,32^. Here, we show that BrtB extends this principle in cyanobacteria by esterifying *in vivo* non-activated FFAs onto bartoloside glycolipids (Figure 2). This study also demonstrates that S06099 tightly controls the levels of B-FAs produced upon FFA supplementation, as cellular levels of B-FA did not increase with increasing concentration of FFA in the medium, while the amount of bartolosides significantly decreased with higher availability of FFA (Figure 3). Despite the saturation levels for B-FAs observed in FFA-supplemented S06099 cells, B-OHs were found to accumulate proportionally to the amount of supplemented FFA (Figure 5). As B-OHs were hypothesized to be hydrolyzed products from B-FAs ^17^, this work shows that FFAs can be released from B-FAs, which suggests the action of a lipase/esterase. Only a limited set of cyanobacterial lipases have been functionally characterized so far. These include LipA (Sll1969), which hydrolyses membrane lipids to FFA ^33^, and the acyl-plastoquinol lipase APL (Sll0482)^34^, both in S6803, and three additional lipases recently identified in *Synechococcus elongatus* PCC 7942 ^35^. This supports the view that lipolytic steps in cyanobacterial lipid turnover remain insufficiently explored. We therefore searched for a lipase candidate genetically associated with the *brt* cluster, which would argue for a dedicated bartoloside ester–hydrolysis module rather than nonspecific background lipase activity. However, no clear lipase candidate was uniquely associated with bartoloside-producing strains.

In addition to B-FAs levels plateauing and bartolosides decreasing while B-OHs increased, membrane lipids PG and SQDG levels also rose upon FFA supplementation (Figure 5). This pattern is consistent with B-FAs acting as transient buffering intermediates rather than a terminal sink for FAs. However, when grown across different temperatures, PG and SQDG fatty-acyl compositions differ, as expected for membrane remodeling ^15,20,36,37^, while the B-FA acyl composition remains largely unchanged (Figure supplement 7). This suggests that the B-FA pool is maintained with a relatively stable acyl profile even when structural membrane lipids adjust. Further experiments will thus be needed to determine whether FFAs released from B-FAs are activated by Aas and redistributed into core membrane lipids, or whether supplemented FFAs that bypass BrtB are directly activated by Aas for incorporation into membrane lipids.

Cyanobacterial responses to exogenous FAs acid are poorly described. Deletion of *aas* in S6803 altered the expression of hundreds of genes, including lipid-associated metabolism ^28^ indicating that extensive FA-related pathways are under transcriptional control. However, and despite a conspicuous remodeling of bartoloside-related metabolites upon FA supplementation, our RNA-seq and quantitative exoproteomics experiments showed no significant global changes in transcript or extracellular protein abundance at the used thresholds (Figure 6 and 7). This contrasts with many heterotrophic bacteria, where exogenous FFA can reprogram FA synthesis and uptake via transcriptional regulators (e.g., the FadR/FabR network in *Escherichia coli* (*E. coli*) coordinating uptake, activation, β-oxidation, and unsaturated FA synthesis) ^38^. We could interpret that, in S06099, the dominant response to incoming FFAs is enzymatic and envelope-proximal, with limited requirement for transcriptional adaptation over the time window examined.

Our observations included a rapid BrtB-dependent esterification of non-activated FFAs which was coupled with BrtB detection at the cell envelope and the absence of a strong response to FFA supplementation. This supports a model in which pre-positioned BrtB intercepts FFA at the cell surface and subsequently routes them into a reversible ester pool. Whether BrtB is a soluble secreted enzyme or an OM-associated protein that is partly recovered extracellularly is still not clear. The presence of an anchor-like domain and detection of the protein in the isolated OM fraction (Figure 7) support the idea that at least part of the BrtB population is displayed at the cell surface. BrtB can then be released into the soluble extracellular fraction via outer-membrane vesicles (OMVs) or other non-specific routes ^41–47^.

Future work will focus on defining the subcellular localization of bartoloside-derived metabolites and on testing their putative structural role at the cell surface ^41^. Extending genetic and imaging tools to non-model cyanobacteria will be key to test these ideas. In summary, our data show that BrtB catalyzes, within minutes, esterification of non-activated FFA onto bartolosides at the cell envelope, generating a reversible B-FA pool that can yield B-OHs and release FFAs. This establishes an activation-independent route for incorporating exogenous FFAs into abundant cyanobacterial metabolites and warrants future work to define metabolite localization and the envelope-level consequences of this chemistry.

## Material and methods

### General experimental procedures

Fatty acids and other substrates used in feeding assays were obtained from Fluorochem (hexanoic acid), Alfa Aesar (octanoic acid), Thermo Fisher Scientific (dodecanoic and palmitic acid), Eurisotop (water ^18^O, 97%) and CDN Isotopes (Octanoic-d_15_-acid, Dodecanoic-d_23_ acid). LC-MS solvents were obtained from Thermo Fisher Scientific, Avantor and Carlo Erba. All solvents were MS-grade.

### Plasmids, strain and culture conditions

The cyanobacterium *Synechocystis salina* LEGE 06099 was obtained from the LEGEcc ^39^ and routinely maintained in liquid culture using Z8 medium supplemented with 25 g L^−1^ sea salt (Tropic Marin), under a light regimen of 16 h light /8 h dark, at 17 °C, 25 °C or 30 °C with agitation on an orbital shaker. For each experiment, cultures were first inoculated using a 3:50 (v/v) inoculum ratio of exponential phase cultures (OD_750_ of ∼0.5). The unicellular cyanobacterium *Synechocystis* sp. PCC 6803 (non-motile and with S-layer ^40^) was routinely maintained in liquid BG11 ^41^ medium, under a light regimen of 16 h light /8 h dark, at 30°C and agitation on an orbital shaker.

### Synthesis of stable isotope ^18^O_2_ hexanoic acid and ^18^O_2_ dodecanoic acid

Stable isotope ^18^O_2_ hexanoic and ^18^O_2_ dodecanoic acid were synthesized based on an adapted version of Murphy et al. ^42^. Labeling with two ^18^O atoms was carried out by simple exchange with H_2_^18^O containing 1 N HCl. Hexanoic and dodecanoic acids were weighed and poured into a 1-mL glass vial to which 200 µL water ^18^O, 97% (Eurisotop) was added. A drop of 37% HCl in deionized H_2_O was added slowly to the solution followed by 20 min sonication. The lead was further sealed with a Teflon-lined tape, the solution constantly stirred within an oil bath at 95 °C for 9 days (hexanoic acid) or three consecutive weeks (dodecanoic acid). The reaction was stopped by evaporation of the solution using a rotary evaporator and further stored at -20 °C for further use. The ^18^O_2_ labelling of the hexanoic acid and dodecanoic acid was confirmed by direct injection into HRESIMS (Figure supplement 1). To directly inject the ^18^O_2_ hexanoic acid and ^18^O_2_ dodecanoic acid, they have been previously solubilized by 10 minutes of sonication in isopropanol containing 5% of deionized H_2_O and 0.75% of acetone.

### Feeding experiments

For supplementation, FA chain length was carefully selected for each assay to balance uptake, solubility, signal, and viability, keeping dosing identical. Importantly, bartoloside lauroyl esters and bartoloside octanoyl esters are the two naturally lowest abundant reported B-FAs in S06099 without supplementation^18^, while bartoloside hexanoyl esters were not detected at all. This provides the basis for a high dynamic range for supplementation assays which makes supplementation-derived products straightforward to quantify. Supplementation of S06099 cultures (in Z8 + sea salt 25 g L^−1^, see above, grown at 25 °C) with ^18^O_2_ hexanoic acid, ^18^O_2_ dodecanoic acid, non-labeled hexanoic acid, non-labeled dodecanoic acid and DMSO was performed in triplicate using a final concentration of 0.75 mM of both labeled and non-labeled hexanoic acid and dodecanoic acid, or no supplementation. The experiments were carried out in 25 mL cultures in Erlenmeyer flasks and cells were harvested after 7 days of growth upon supplementation by centrifugation at 4500 × *g*, 10 min, 4 °C, rinsed with deionized H_2_O and centrifuged again. The resulting biomass pellets were lyophilized prior to extraction with CH_2_Cl_2_/MeOH (2:1, v/v) at room temperature. Crude extracts were analyzed at a concentration of 1 mg mL^-1^ dissolved in methanol, by LC-HRESIMS and LC-HRESIMS/MS.

Supplementation of S06099 cultures (in Z8 + sea salt 25 g L^−1^, see above, grown at 25 °C) with ^D^-octanoic acid, ^D^-dodecanoic acid, non-labeled octanoic acid, non-labeled dodecanoic acid and DMSO was performed in triplicate using a final concentration of 0.01 mM, 0.05 mM and 0.5 mM of both labeled and non-labeled hexanoic acid and dodecanoic acid in 1% of final volume DMSO, or only 1% final volume DMSO. The dose-response experiments were carried out in 25 mL cultures in Erlenmeyer flasks and cells were harvested after 24 h and 7 days of growth upon supplementation by centrifugation at 4500 × *g*, 10 min, 4 °C, rinsed with deionized H_2_O and centrifuged again. The resulting biomass pellets were lyophilized prior to extraction with CH_2_Cl_2_/MeOH (2:1, v/v) at room temperature. Crude extracts were analyzed at a concentration of 1 mg mL^-1^ dissolved in methanol, by LC-HRESIMS and LC-HRESIMS/MS.

### LC-HRESIMS analysis

For LC-HRESIMS analyses, separation was performed in a Luna C18 column (100 × 3 mm, 3 μm, 100 Å, Phenomenex). Mixtures of MeOH/ H_2_O 1:1 (v/v) with 0.1% formic acid (eluent A) and IPA with 0.1% formic acid (eluent B) were used as mobile phase, with a flow rate of 0.4 mL min^−1^, 10 μL of a 1 mg mL^−1^ solution were injected and separated using a gradient from at 9:1 to 1:4 eluent A/eluent B in 3 min and held for 28 min before returning to initial conditions.

Analysis of ^18^O_2_ labelling of the hexanoic acid and dodecanoic acid was carried out by injecting 10 μL of the labeled fatty acids. Separation involved an isocratic step of 9:1 eluent A/eluent B over 2 min, followed by a linear gradient to 7:13 eluent A/eluent B over 3 min and held for 10 min, followed by a linear gradient to 3:17 eluent A/eluent B over 3 min and held for 8 min before returning to the initial conditions (Figure supplement 1).

LC-HRESIMS (HCD, Higher energy collisional dissociation) analyses were performed on an UltiMate 3000 UHPLC (Thermo Fisher Scientific) system composed of a LPG-3400SD pump, WPS-3000SL autosampler and VWD-3100 UV/VIS detector coupled to a Q Exactive Focus Hybrid Quadrupole-Orbitrap Mass Spectrometer controlled by Q Exactive Focus Tune 2.9 and Xcalibur 4.1 (Thermo Fisher Scientific). LC-HRESIMS data were obtained in Full Scan mode, with a capillary voltage of HESI set to −3.8 kV and the capillary temperature to 300 °C. Sheath gas flow rate was set to 35 units.

### Genome assembly and proteome generation

The S06099 genome assembly (WGS accession JADEWK000000000; currently processing at NCBI) was used as the reference for all omics analyses. Protein-coding genes were predicted with Prodigal, yielding 3372 predicted proteins, and functional annotation was assigned with eggNOG-mapper v2 (DIAMOND mode). The resulting predicted proteome FASTA was used for transcriptomic and proteomics library prediction and database searches.

### S06099 RNA extraction and sequencing

Experimental setup is elaborated in Figure 3A. Samples for total RNA and targeted metabolomics analysis were taken at 30 min and 6 h upon FA supplementation. Bacterial cell pellets were obtained from the same culture for both metabolites and RNA extraction. For each replicate, 15 mL of culture for metabolites extraction was centrifuge at 4 500 × *g* for 10 min at 4 °C then rinsed with deionized H_2_O and centrifuged again for 10 min; and 100 mL of culture for RNA extraction was centrifuge at 3,000 × *g* for 10 min followed by 3,000 × g, 5 min, all at 4 °C. Cell pellets were suspended in 1 mL RNA*later* Stabilization buffer (Thermo Fisher Scientific) and incubated at room temperature for 5 min, followed by centrifugation at 3,000 × *g*, for 5 min. Cells were stored at −80 °C until further analysis. Then, cells were flash frozen in liquid nitrogen and disrupted by mortar and pestle. RNA extraction was carried out using the PureLink^TM^ RNA Mini Kit (Thermo Fisher Scientific) according to the manufacturer’s protocol. DNA was removed using RapidOut DNA Removal Kit (Thermo Fisher Scientific); RNA-Seq library preparation and subsequent Illumina sequencing (150 bp paired-end) were performed by Novogene.

### Transcriptome analysis

Raw reads (364.4 million) were quality checked using FastQC (www.bioinformatics.babraham.ac.uk/projects/fastqc), and quality trimmed using BBDuk from BBMap ^43,44^ (qtrim=rl ; trimq=20). Reads were mapped to S06099 genome assembly (NCBI GenBank under the WGS accession JADEWK000000000, currently processing; public release upon acceptance) using Bowtie2 ^45^ and reads were counted on genes using FeatureCounts ^46^. Gene expression was quantified using DESeq2 ^47^. Genes were considered DE when |log2 fold change| ≥ 1 and adjusted p value ≤ 0.05 (Benjamini–Hochberg). PCA and similarity between samples were calculated using normalized gene expression values and visualized using ggplot RStudio 3.5.0. Volcano plots of gene expression were generated using the EnhancedVolcano R package ^48^. Transcriptomic data were submitted to ArrayExpress/BioStudies^29,49^ via Annotare, with sequencing reads brokered to ENA available through ENA series accession number E-MTAB-16709.

### Protein structure prediction and visualization

The three-dimensional structure of BrtB was predicted using the AlphaFold Protein Structure Database (model AF-A0A1B3Z488-F1-v4). Structural visualization and annotation were performed in PyMOL 3.1.3 (Schrödinger, LLC). Domain regions and specific sequence features were highlighted based on residue numbering, including the β-propeller core, the C-terminal region and proline-enriched segments. Images were exported as high-resolution PNG files for figure assembly.

### Preparation and analysis of S6803 and S06099 outer membrane, cytoplasmic and cell free extracellular proteins

The outer membranes of S6803 and S06099 were isolated according to the method used in Cardoso et al. ^50^. The cells were grown in 100 mL of medium until the cultures reached an OD_750_ of ∼ 0.8-1, under the same conditions as described (strain and culture condition) at 30 °C. In brief, cells were collected by centrifugation at 4400× *g* for 10 min at room temperature and the corresponding pellet was washed in 1 mL of buffer O (10 mM Tris-HCl, pH 7.45). The mixture was then transferred to a 1.5 mL microfuge tube and centrifuged at 16,100× *g* for 2 min at 4 °C. Cells were suspended in 400 μL of buffer O and lysed by sonication (Bandelin Sonoplus (50% duty cycle)) in 6 cycles of 15 sec power, 15 sec on ice. Then, cell lysates were centrifuged for 15 sec at 16,100× *g* (4 °C) to remove unbroken cells and heavy cellular debris, and the supernatant was further centrifuged for 30 min at 16,100× g (4 °C). The resulting pellet was washed in 200 μL of buffer O and centrifuged once again for 30 min under the same conditions. The supernatant containing cytoplasmic proteins was stored at −80 °C. Next, the pellet was suspended in 200 μL buffer O and mixed with 250 μL of buffer O containing 2% (v/v) N-Lauroylsarcosine. The mixture was incubated for 30 min with agitation in a Fisherbrand^TM^ Nutating Mixer (Thermo Fisher Scientific) at room temperature. Outer membranes were then collected by centrifugation (45 min at 16,100× *g* (4 °C)), and washed twice in 400 μL of buffer O. Finally, isolated OM proteins were resuspended in 50 μL of buffer O and stored at −80 °C.

After cell harvesting, conditioned medium was filtered through 0.2 μm pore-size filters and concentrated down to 50 µL by ultrafiltration with Amicon Ultra-4 Centrifugal Filter units (Millipore) with a nominal molecular weight limit of 3 kDa ^50^ and stored at −80 °C. Proteins were separated by electrophoresis on homogeneous 12% (w/v) SDS-polyacrylamide gels. Peptide band detection was performed with Bio-Safe Coomassie G-250 Stain (Bio-Rad).

BrtB expressed in *E. coli* with a N-terminal histidine tag (His-BrtB) was obtained as described in Reis et al. ^18^.

To analyze the exoproteome of S06099 by the EXCRETE workflow ^29^, cells were grown in the same conditions as described above, with or without supplementation of 0.4 mM of hexadecenoic acid for 24 h, in 4 replicates. Palmitic acid was chosen as it is prevalent in cyanobacterial lipids ^30^ and bartolosides palmitate esters are the most abundant B-FAs in S06099, making C16:0 the substrate that most closely reflects the native bartoloside pool, and thus the most suitable probe for BrtB localization. Cells present in 50 mL of each cell culture were pelleted at 4400 × *g* for 10 min at room temperature, extracellular media filtered, and samples stored at -80 °C until preparation. Subsequent sample preparation for exoproteomic analysis was done according to previously published protocols ^29,49^ except that the sample amount used was determined by volume (900 µL per sample) rather than protein amount due to low protein concentrations (< 5 µg mL^-1^).

### Protein identification by mass spectrometry

Selected polyacrylamide gel bands present in S06099 protein fractions and absent from S6803 were identified by mass spectrometry at the Proteomic Unit, i3S, University of Porto, Portugal. Gel bands were washed twice with 50% acetonitrile (ACN) in 50 mM triethylammonium bicarbonate (TEAB) with shaking at 1500 rpm for 5 min and further treated with ACN twice. Next, proteins were reduced with 25 mM dithiothreitol (DTT) for 20 min at 56 °C and alkylated with 55 mM iodoacetamide (IAA) for 20 min at room temperature in the dark, followed by the same wash procedure. Proteins were then digested with trypsin (240 ng) in 50 mM TEAB/0.01% (w/v) surfactant for 60 min at 50 °C. Peptide gel extraction was performed with 2.5% trifluoroacetic acid (TFA) followed by 50% ACN, 1% TFA. Then, samples were dried, suspended in 10 μL 0.1% TFA and cleaned by C18 reverse phase chromatography following the manufacturer’s instructions (ZipTip, Merck, Darmstadt, Germany). Sample protein identification and quantification was performed with a 120 min chromatographic separation run by nano liquid chromatography mass spectrometry (nano LC-MS/MS) equipped with a Field Asymmetric Ion Mobility Spectrometry - FAIMS interface. This equipment is composed of a Vanquish Neo liquid chromatography system coupled to an Eclipse Tribrid Quadrupole, Orbitrap, Ion Trap mass spectrometer (Thermo Scientific, San Jose, CA). The specific MS parameters were: MS maximum injection time, auto; dd settings: normalized AGC target 100%, absolute AGC value 1.0 × 10^4^, intensity threshold 5.0 × 10^3^, and dynamic exclusion 30 s. Data acquisition was controlled by Tune 4.0 software (Thermo Scientific, San Jose, CA, USA). S06099 assembled proteome was considered for protein identification. Protein identification was performed with the Proteome Discoverer software (v3.1.1.93, Thermo Scientific).

Purified and desalted peptides were separated using a 30-min gradient and analyzed using the diaPASEF acquisition mode ^51^ on a trapped ion mobility spectrometry (TIMS) quadrupole time-of-flight mass spectrometer as described in Russo, Schneidmadel and Zedler, 2025 ^52^. DIA raw files were analyzed using DIA-NN version 2.1 ^53^ with default settings in library-free mode. A predicted S06099 proteome (3372 proteins), generated from the genome assembly by Prodigal gene prediction (see Genome assembly and annotation), was used for library prediction. Deep learning-based prediction of spectra, retention times, and ion mobilities was enabled. Trypsin/P was specified as the protease, allowing up to two missed cleavages. Methionine oxidation (Ox[M]) was set as a fixed modification. Match-between-runs was enabled across all samples. A precursor-level FDR of 1% was applied. A protein group was considered identified when it was present in at least 70% of the replicates with a minimum of three replicates. Prediction of protein function and location was done using EggNOG 5.0 ^54^and DeepLocPro 1.0 ^55^.

### Transmission electron microscopy

Cells in 5 mL of each cell culture grown until OD_750_ ∼ 0.8-1, with or without supplementation of 0.4 mM of palmitic acid for 24 h, in duplicate were collected for ultrastructural analysis. Cells were fixed overnight at 4 °C in a solution containing 2% formaldehyde and 2.5% glutaraldehyde in 0.1 M sodium cacodylate buffer. Then, cells were washed with 0.1 M sodium cacodylate buffer, embedded in Histogel™, and post-fixed for 2 hours in 2% osmium tetroxide in 0.1 M sodium cacodylate buffer. Afterward, cells were washed in water, stained with aqueous 1% uranyl acetate for 30 minutes, dehydrated, and embedded in Embed-812 resin. Ultra-thin sections (70 nm thick) were cut using an RMC Ultramicrotome with Diatome diamond knives, mounted on 200 mesh copper grids, and stained with uranyl and 3% lead citrate for 5 minutes each. Imaging was done using a JEOL JEM 1400 transmission electron microscope, and digital images were captured with a PHURONA CCD camera. Transmission electron microscopy was performed at the Histology and Electron Microscopy core facility at i3S, University of Porto, Portugal, with the assistance of Sofia Pacheco and Rui Fernandes. Cell size and the distance between the outer membrane and the S-layer were measured in Fiji (ImageJ v2.12.0). Two biological replicates of S06099 were analyzed; for each replicate, 15 micrographs were quantified, and for the S-layer to outer-membrane distance, 10 positions per cell were measured.

### Knockout attempt of BrtB in S06099

A non-replicative brtB knock out plasmid was assembled in pBluescript II using upstream (B1) and downstream (B2) homology arms amplified from S06099 gDNA (523 and 531 base pair, respectively). The arms were amplified with Phusion polymerase in GC buffer supplemented with DMSO (annealing 64 °C) using primers: B1_Forward CTCGAGGGGCAATTTTATCCACTGTC); B1_Reverse CTGCAGGAATAGCGATTAACGATTACGG); B2_Forward CTGCAGTTGAACGGTGGTGCTTCC); B2_Reverse GGATCCGTCGGTGTAATTAAACCAAGCATAAG), which introduce restriction sites XhoI/PstI and PstI/BamHI. Purified amplicons were digested (B1: XhoI/PstI; B2: PstI/BamHI) and ligated into PstI-linearized pBluescript II at a 1:3 insert:vector ratio using T4 DNA ligase. *E. coli* transformants were screened by restriction analysis (XhoI/PstI and XhoI/BamHI) and one clone was sequence-verified. To insert the kanamycin resistance cassette between the two arms, the intermediate plasmid was digested with PstI and dephosphorylated (alkaline phosphatase), ligated with the PstI-flanked kanamycin cassette at a 1:3 insert:vector ratio, and screened by restriction analysis (PstI and XhoI) and Sanger sequencing (B1/B2 primers, kanamycin cassette reverse primer, and M13 reverse). For natural transformation, mid-exponential S06099 cultures (OD ∼0.5) were mixed with 4 to 10 μg plasmid DNA; washed or non-washed cell preparations were both tested; and plated either directly on agar or on nitrocellulose membranes Z8 supplemented with 25 g L^−1^ sea salt. After recovery on non-selective plates, cells were transferred to selective plates containing 15 or 30 μg mL-1 kanamycin and incubated at 23 °C, 25 °C or 30 °C. For electroporation, cultures grown to OD ∼0.65 were washed and concentrated, mixed with 1 μg of plasmid DNA, electroporated, and processed through the same recovery and selection steps. Putative kanamycin-resistant colonies were restreaked on higher kanamycin concentrations (50 to 100 μg mL-1) and screened by colony PCR using brtB- and kanamycin-specific primers (Q5; 98 °C 3 min; 30 to 35 cycles of 98 °C 10 s, 67 °C 30 s, 72 °C 40 s; final extension 72 °C 2 min). Across repeated rounds of selection, segregation and PCR screening, no stable kanamycin-resistant isolate consistent with a brtB replacement was obtained. In parallel, plasmid transfer into S06099 was tested by conjugation using the replicative vector pEF07 (pSEVA251::sfGFP; RSF1010 origin; Ptrc.x.lacO promoter). Conjugation followed a triparental mating format using *E. coli* HB101 carrying pEF07 and the helper plasmid pRL623, together with an *E. coli* strain carrying the conjugative plasmid pRL443. Mating mixtures were incubated on non-selective BG11 agar supplemented with 25 g L^−1^ sea salt medium plates at 30 °C for 24 h before recovery (cells were resuspended in 750 μL growth medium, centrifuged at 4500 × g for 5 to 8 min and concentrated to ∼300 μL) and replating on selective agar (250 μL onto 30 μg mL-1 kanamycin and 50 μL onto 15 μg mL-1 kanamycin). Plates were incubated at 30 °C under a 14 h light / 10 h dark regime and monitored for at least two weeks by visual inspection and fluorescence microscopy, but no stable GFP-expressing exconjugants were recovered.

### Statistics and reproducibility

All experiments were conducted with three to four biological replicates unless stated otherwise. Mean values are reported and individual replicate points are shown in the figures. Details on sample sizes, processing steps, and normalization procedures are provided in the respective subsections and figure legends. For FA supplementation assays, replicate-specific sums were computed at the metabolite-family level, and differences across feeding conditions were evaluated by one-way ANOVA followed by Tukey’s post-hoc test for all pairwise comparisons. Tukey-adjusted p-values are reported. For targeted metabolomics data accompanying the transcriptomics experiment, replicate-level family sums were analyzed using a linear model including Feeding, Time_point, and their interaction (Feeding x Time_point) on log1p-transformed values. Post-hoc contrasts were estimated from model marginal means (emmeans) to (i) compare C12 versus DMSO within each time point, and (ii) compare 6 h versus 30 min within the C12 condition; *p*-values were adjusted across families using the Benjamini-Hochberg procedure within each contrast set. Only metabolites detected in at least two replicates were included. For RNA-seq analyses, normalization, dispersion estimation and differential expression testing followed the DESeq2 workflow (details in the RNA-seq Methods). Temperature-dependent comparisons were computed per replicate long-chain fractions (≥ C34 for PG/SQDG; ≥ C14 for B-FAs) and unsaturation indices (ΣProportion × double bonds / 100) from the species-resolved data. Temperature effects (17 °C vs 30 °C) on these indices were tested separately for each time point and lipid class using Welch’s t-test with Benjamini–Hochberg correction (rstatix in R). All statistical analyses were performed in R (version 4.4.2) using base functions, ggplot2 (3.5.1), and DESeq2 (1.50.2).

## Acknowledgements

The authors acknowledge the support of the i3S Scientific Platform HEMS, member of the national infrastructure PPBI – Portuguese Platform of Bioimaging (PPBI-POCI-01-0145-FEDER-022122). We thank the Microbial Diversity course (2024 and 2025) from the Marine Biological Laboratory and the Engel laboratory (Biozentrum, University of Basel) for critical discussions that informed the imaging aspects of this work.

## Data availability

Whole-genome shotgun (WGS) sequencing data for *Synechocystis salina* LEGE 06099 have been submitted to NCBI GenBank under the WGS accession JADEWK000000000 (currently processing; public release upon acceptance).

RNA-seq data have been deposited in ENA under accession E-MTAB-16709.

The mass spectrometry exoproteomics data have been deposited to the ProteomeXchange Consortium via the PRIDE ^56^partner repository with the dataset identifier PXD074249. The data is available for review using the username: and password: 3RjbBMINalgv.

## Author details

Amaranta kahn, Conceptualization, Methodology, Formal analysis, Investigation, Data curation, Writing-original draft, Writing-review & editing, Visualization, Supervision, Funding acquisition; Cristina I.F Sousa, Methodology, Formal analysis, Investigation, Data curation, Visualization; João P. A. Reis, Methodology, Formal analysis, Investigation; David A. Russo, Methodology, Formal analysis, Investigation, Data curation, Writing-review & editing, Visualization; Adriana Rego, Methodology, Formal analysis; Ricardo J.C.V Queirós, Investigation; Marine Cuau, Investigation; Julie A. Z. Zedler, Methodology; Sandra A.C. Figueiredo, Methodology, Formal analysis, Investigation, Data curation; Paulo Oliveira, Conceptualization, Methodology, Formal analysis, Investigation, Writing-review & editing; Pedro N. Leão, Conceptualization; Methodology; Validation; Formal analysis; Resources; Writing–review & editing; Visualization; Supervision; Project administration; Funding acquisition.

## References

1. Wältermann, M., & Steinbüchel, A. (2005). Neutral lipid bodies in prokaryotes: Recent insights into structure, formation, and relationship to eukaryotic lipid depots. Journal of Bacteriology, 187(11), 3607–3619. 10.1128/JB.187.11.3607-3619.2005

2. Mathiowetz, A. J., & Olzmann, J. A. (2024). Lipid droplets and cellular lipid flux. In Nature Cell Biology (Vol. 26, Number 3). 10.1038/s41556-024-01364-4

3. Resh, M. D. (2016). Fatty acylation of proteins: The long and the short of it. Progress in Lipid Research, 63, 120–131. 10.1016/j.plipres.2016.05.002

4. Resh, M. D. (2006). Trafficking and signaling by fatty-acylated and prenylated proteins. In Nature Chemical Biology (Vol. 2, Number 11). 10.1038/nchembio834

5. Justice, I., Kiesel, P., Safronova, N., von Appen, A., & Saenz, J. P. (2024). A tuneable minimal cell membrane reveals that two lipid species suffice for life. Nature Communications, 15(1), 9679. 10.1038/s41467-024-53975-y

6. Huang, H., Wang, C., Chang, S., Cui, T., Xu, Y., Huang, M., Zhang, H., Zhou, C., Zhang, X., & Feng, Y. (2025). Structure and catalytic mechanism of exogenous fatty acid recycling by AasS, a versatile acyl-ACP synthetase. Nature Structural and Molecular Biology, 32(5). 10.1038/s41594-024-01464-7

7. Beld, J., Lee, D. J., & Burkart, M. D. (2015). Fatty acid biosynthesis revisited: Structure elucidation and metabolic engineering. In Molecular BioSystems (Vol. 11, Number 1). 10.1039/c4mb00443d

8. Yao, J., & Rock, C. O. (2017). Exogenous fatty acid metabolism in bacteria. In Biochimie. 10.1016/j.biochi.2017.06.015

9. Kahn, A., Oliveira, P., Cuau, M., & Leão, P. N. (2023). Incorporation, fate, and turnover of free fatty acids in cyanobacteria. FEMS Microbiology Reviews, 47(2), 1–19. 10.1093/femsre/fuad015

10. Currie, M. F., Persaud, D. M., Rana, N. K., Platt, A. J., Beld, J., & Jaremko, K. L. (2020). Synthesis of an acyl-acyl carrier protein synthetase inhibitor to study fatty acid recycling. Scientific Reports. 10.1038/s41598-020-74731-4

11. von Berlepsch, S., Kunz, H.-H., Brodesser, S., Fink, P., Marin, K., Flügge, U.-I., & Gierth, M. (2012). The Acyl-Acyl Carrier Protein Synthetase from *Synechocystis* sp. PCC 6803 Mediates Fatty Acid Import. Plant Physiology, 159(2), 606–617. 10.1104/pp.112.195263

12. Kaczmarzyk, D., & Fulda, M. (2010). Fatty acid activation in cyanobacteria mediated by acyl-acyl carrier protein synthetase enables fatty acid recycling. Plant Physiology, 152(3), 1598–1610. 10.1104/pp.109.148007

13. Eungrasamee, K., Miao, R., Incharoensakdi, A., Lindblad, P., & Jantaro, S. (2019). Improved lipid production via fatty acid biosynthesis and free fatty acid recycling in engineered *Synechocystis* sp. PCC 6803. Biotechnology for Biofuels, 12(1). 10.1186/s13068-018-1349-8

14. Leão, P. N., Nakamura, H., Costa, M., Pereira, A. R., Martins, R., Vasconcelos, V., Gerwick, W. H., & Balskus, E. P. (2015). Biosynthesis-Assisted Structural Elucidation of the Bartolosides, Chlorinated Aromatic Glycolipids from Cyanobacteria. Angewandte Chemie - International Edition. 10.1002/anie.201503186

15. Figueiredo, S. A. C., Preto, M., Moreira, G., Martins, T. P., Abt, K., Melo, A., Vasconcelos, V. M., & Leão, P. N. (2021). Discovery of cyanobacterial natural products containing fatty acid residues. Angewandte Chemie - International Edition.

16. Afonso, T. B., Costa, M. S., Rezende De Castro, R., Freitas, S., Silva, A., Schneider, M. P. C., Martins, R., & Leão, P. N. (2016). Bartolosides E-K from a Marine coccoid cyanobacterium. Journal of Natural Products. 10.1021/acs.jnatprod.6b00351

17. Reis, J. P. A., Freitas, S., Procházková, T., & Leão, P. N. (2024). Expanding the Diversity of the Cyanobacterial Dialkylresorcinol Bartoloside Family. Journal of Natural Products. 10.1021/acs.jnatprod.4c00832

18. Reis, J. P. A., Figueiredo, S. A. C., Sousa, M. L., & Leão, P. N. (2020). BrtB is an O-alkylating enzyme that generates fatty acid-bartoloside esters. Nature Communications. 10.1038/s41467-020-15302-z

19. Wada, H., & Murata, N. (1998). Membrane Lipids in Cyanobacteria. In Lipids in Photosynthesis: Structure, Function and Genetics (pp. 65–81). Kluwer Academic Publishers. 10.1007/0-306-48087-5_4

20. Wada, H., & Murata, N. (1990). Temperature-induced changes in the fatty acid composition of the cyanobacterium, *Synechocystis* PCC6803. Plant Physiology, 92(4), 1062–1069. 10.1104/pp.92.4.1062

21. Jaroensuk, J., Abraham, J. P., Zuniga, B. E., Shepard, H. S., Wei, M., Williams, R., Morley, S. A., Lingwan, M., Zhou, J., Jindra, M. A., Jyoti, P., Wang, B., May, J. C., McLean, J. A., Young, J. D., Pfleger, B. F., & Allen, D. K. (2026). Disruption of acyl-acyl carrier protein (acyl-ACP) synthetase in cyanobacteria impairs lipid remodeling as revealed by acyl-ACP measurements. Metabolic Engineering, 94. 10.1016/j.ymben.2025.11.004

22. Sinetova, M. A., & Los, D. A. (2016). Systemic analysis of stress transcriptomics of *Synechocystis* reveals common stress genes and their universal triggers. Molecular BioSystems, 12(11). 10.1039/c6mb00551a

23. Lee, S., Ryu, J. Y., Soo, Y. K., Jeon, J. H., Ji, Y. S., Cho, H. T., Choi, S. B., Choi, D., De Marsac, N. T., & Park, Y. Il. (2007). Transcriptional regulation of the respiratory genes in the cyanobacterium *Synechocystis* sp. PCC 6803 during the early response to glucose feeding. Plant Physiology, 145(3). 10.1104/pp.107.105023

24. Zhang, Z., Pendse, N. D., Phillips, K. N., Cotner, J. B., & Khodursky, A. (2008). Gene expression patterns of sulfur starvation in *Synechocystis* sp. PCC 6803. BMC Genomics, 9. 10.1186/1471-2164-9-344

25. Osanai, T., Kanesaki, Y., Nakano, T., Takahashi, H., Asayama, M., Shirai, M., Kanehisa, M., Suzuki, I., Murata, N., & Tanaka, K. (2005). Positive regulation of sugar catabolic pathways in the cyanobacterium *Synechocystis* sp. PCC 6803 by the group 2 σ factor SigE. Journal of Biological Chemistry, 280(35). 10.1074/jbc.M505043200

26. Liu, Y., Tsinoremas, N. F., Johnson, C. H., Lebedeva, N. V., Golden, S. S., Ishiura, M., & Kondo, T. (1995). Circadian orchestration of gene expression in cyanobacteria. Genes and Development, 9(12). 10.1101/gad.9.12.1469

27. Johnson, C. H., Mori, T., & Xu, Y. (2008). A Cyanobacterial Circadian Clockwork. In Current Biology (Vol. 18, Number 17). 10.1016/j.cub.2008.07.012

28. Gao, Q., Tan, X., & Lu, X. (2013). Enzymatic and physiological characterization of fatty acid activation in *Synechocystis* sp. PCC6803. Journal of Basic Microbiology, n/a-n/a. 10.1002/jobm.201200228

29. Russo, D. A., Oliinyk, D., Pohnert, G., Meier, F., & Zedler, J. A. Z. (2024). EXCRETE workflow enables deep proteomics of the microbial extracellular environment. Communications Biology, 7(1), 1189. 10.1038/s42003-024-06910-2

30. Hewelt-Belka, W., Kot-Wasik, Á., Tamagnini, P., & Oliveira, P. (2020). Untargeted Lipidomics Analysis of the Cyanobacterium *Synechocystis* sp. PCC 6803: Lipid Composition Variation in Response to Alternative Cultivation Setups and to Gene Deletion. International Journal of Molecular Sciences, 21(23), 8883. 10.3390/ijms21238883

31. Yamashita, A., Hayashi, Y., Matsumoto, N., Nemoto-Sasaki, Y., Koizumi, T., Inagaki, Y., Oka, S., Tanikawa, T., & Sugiura, T. (2017). Coenzyme-a-independent transacylation system; possible involvement of phospholipase a2 in transacylation. In Biology (Vol. 6, Number 2). 10.3390/biology6020023

32. Dahlqvist, A., Ståhl, U., Lenman, M., Banas, A., Lee, M., Sandager, L., Ronne, H., & Stymne, S. (2000). Phospholipid:diacylglycerol acyltransferase: An enzyme that catalyzes the acyl-CoA-independent formation of triacylglycerol in yeast and plants. Proceedings of the National Academy of Sciences of the United States of America, 97(12). 10.1073/pnas.120067297

33. Kojima, K., Matsumoto, U., Keta, S., Nakahigashi, K., Ikeda, K., Takatani, N., Omata, T., & Aichi, M. (2022). High-Light-Induced Stress Activates Lipid Deacylation at the Sn-2 Position in the Cyanobacterium *Synechocystis* Sp. PCC 6803. Plant and Cell Physiology, 63(1), 82–91. 10.1093/pcp/pcab147

34. Jimbo, H., Torii, M., Fujino, Y., Tanase, Y., Kurima, K., Sato, N., & Wada, H. (2024). Acyl-turnover of acylplastoquinol enhances recovery of photodamaged PSII in *Synechocystis*. Plant Journal. 10.1111/tpj.17051

35. Takatani, N., Uenosono, M., Senoo, Y., Ikeda, K., Aichi, M., & Omata, T. (2025). A galactolipase activated by high light helps cells acclimate to stress in cyanobacteria. Plant Physiology, 197(4). 10.1093/plphys/kiaf130

36. Nanjo, Y., Mizusawa, N., Wada, H., Slabas, A. R., Hayashi, H., & Nishiyama, Y. (2010). Synthesis of fatty acids de novo is required for photosynthetic acclimation of *Synechocystis* sp. PCC 6803 to high temperature. Biochimica et Biophysica Acta - Bioenergetics, 1797(8), 1483–1490. 10.1016/j.bbabio.2010.03.014

37. Sakamoto, T., & Bryant, D. A. (2002). Synergistic effect of high-light and low temperature on cell growth of the Δ12 fatty acid desaturase mutant in *Synechococcus* sp. PCC 7002. Photosynthesis Research. 10.1023/A:1019820813257

38. Waters, J. K., & Eijkelkamp, B. A. (2024). Bacterial acquisition of host fatty acids has far-reaching implications on virulence. Microbiology and Molecular Biology Reviews, 88(4). 10.1128/mmbr.00126-24

39. Ramos, V., Morais, J., Castelo-Branco, R., Pinheiro, Â., Martins, J., Regueiras, A., Pereira, A. L., Lopes, V. R., Frazão, B., Gomes, D., Moreira, C., Costa, M. S., Brûle, S., Faustino, S., Martins, R., Saker, M., Osswald, J., Leão, P. N., & Vasconcelos, V. M. (2018). Cyanobacterial diversity held in microbial biological resource centers as a biotechnological asset: the case study of the newly established LEGE culture collection. Journal of Applied Phycology. 10.1007/s10811-017-1369-y

40. Oliveira, P., Martins, N. M., Santos, M., Pinto, F., Büttel, Z., Couto, N. A. S., Wright, P. C., & Tamagnini, P. (2016). The versatile TolC-like Slr1270 in the cyanobacterium *S ynechocystis* sp. PCC 6803. Environmental Microbiology, 18(2), 486–502. 10.1111/1462-2920.13172

41. Rippka, R., Deruelles, J., & Waterbury, J. B. (1979). Generic assignments, strain histories and properties of pure cultures of cyanobacteria. Journal of General Microbiology, 111(1). 10.1099/00221287-111-1-1

42. Murphy, R. C., & Clay, K. L. (1990). Preparation of labeled molecules by exchange with oxygen-18 water. Methods in Enzymology, 193(C). 10.1016/0076-6879(90)93425-K

43. Bushnell, B. (2021). BBDuk Guide - DOE Joint Genome Institute. DOE Joint Genome Institute.

44. Bushnell, B. (2015). BBMap. Https://Sourceforge.Net/Projects/Bbmap/.

45. Langmead, B., & Salzberg, S. L. (2012). Fast gapped-read alignment with Bowtie 2. Nature Methods, 9(4). 10.1038/nmeth.1923

46. Liao, Y., Smyth, G. K., & Shi, W. (2014). FeatureCounts: An efficient general purpose program for assigning sequence reads to genomic features. Bioinformatics, 30(7). 10.1093/bioinformatics/btt656

47. Love, M. I., Huber, W., & Anders, S. (2014). Moderated estimation of fold change and dispersion for RNA-seq data with DESeq2. Genome Biology, 15(12). 10.1186/s13059-014-0550-8

48. Blighe, K., Rana, S., & Lewis, M. (2024). EnhancedVolcano: Publication-ready volcano plots with enhanced colouring and labeling. R package version 1.20.0. Https://Bioconductor.Org/Packages/EnhancedVolcano.

49. Teikari, J. E., Russo, D. A., Heuser, M., Baumann, O., Zedler, J. A. Z., Liaimer, A., & Dittmann, E. (2025). Competition and interdependence define interactions of Nostoc sp. and Agrobacterium sp. under inorganic carbon limitation. Npj Biofilms and Microbiomes, 11(1). 10.1038/s41522-025-00675-0

50. Cardoso, D., Lima, S., Matinha-Cardoso, J., Tamagnini, P., & Oliveira, P. (2021). The role of outer membrane protein(S) harboring slh/oprb-domains in extracellular vesicles’ production in *synechocystis* sp. pcc 6803. Plants, 10(12). 10.3390/plants10122757

51. Meier, F., Brunner, A. D., Frank, M., Ha, A., Bludau, I., Voytik, E., Kaspar-Schoenefeld, S., Lubeck, M., Raether, O., Bache, N., Aebersold, R., Collins, B. C., Röst, H. L., & Mann, M. (2020). diaPASEF: parallel accumulation–serial fragmentation combined with data-independent acquisition. Nature Methods, 17(12). 10.1038/s41592-020-00998-0

52. Russo, D. A., Schneidmadel, F. R., & Zedler, J. A. Z. (2025). Library-free data-independent acquisition mass spectrometry enables comprehensive coverage of the cyanobacterial proteome. Plant Physiology, 199(1). 10.1093/plphys/kiaf334

53. Demichev, V., Messner, C. B., Vernardis, S. I., Lilley, K. S., & Ralser, M. (2020). DIA-NN: neural networks and interference correction enable deep proteome coverage in high throughput. Nature Methods, 17(1). 10.1038/s41592-019-0638-x

54. Huerta-Cepas, J., Szklarczyk, D., Heller, D., Hernández-Plaza, A., Forslund, S. K., Cook, H., Mende, D. R., Letunic, I., Rattei, T., Jensen, L. J., Von Mering, C., & Bork, P. (2019). EggNOG 5.0: A hierarchical, functionally and phylogenetically annotated orthology resource based on 5090 organisms and 2502 viruses. Nucleic Acids Research, 47(D1). 10.1093/nar/gky1085

55. Moreno, J., Nielsen, H., Winther, O., & Teufel, F. (2024). Predicting the subcellular location of prokaryotic proteins with DeepLocPro. BioRxiv, (Dl).

56. Perez-Riverol, Y., Bandla, C., Kundu, D. J., Kamatchinathan, S., Bai, J., Hewapathirana, S., John, N. S., Prakash, A., Walzer, M., Wang, S., & Vizcaíno, J. A. (2025). The PRIDE database at 20 years: 2025 update. Nucleic Acids Research, 53(D1). 10.1093/nar/gkae1011

