## Supplementary material for "Activation-independent capture of free fatty acids at the bacterial cell envelope by BrtB": Figure supplements

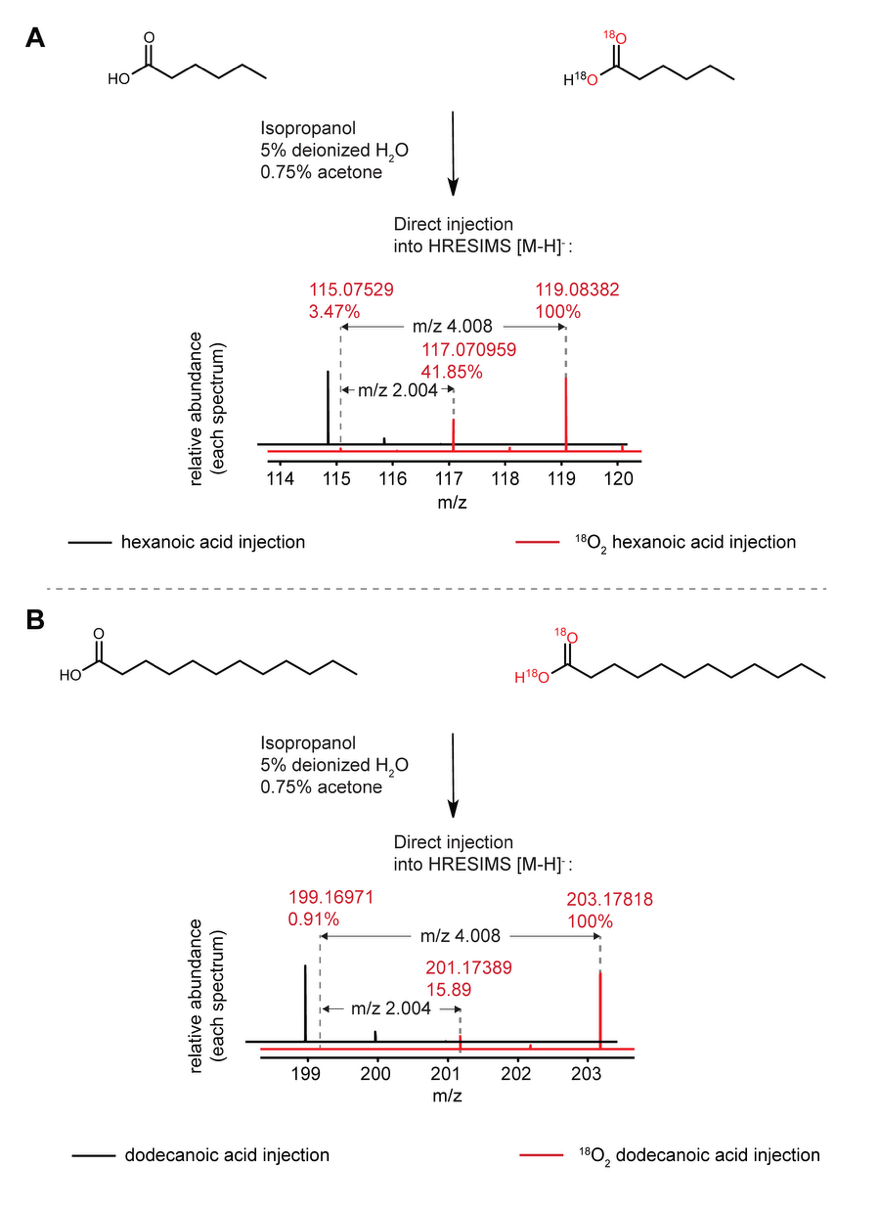

**Figure supplement 1.** **Confirmation and quantification of ¹⁸O₂ labeling of hexanoic and dodecanoic acids by direct injection HRESIMS.** Stable isotope ¹⁸O₂ hexanoic acid and ¹⁸O₂ dodecanoic acid were generated by oxygen exchange in H₂¹⁸O under acidic conditions (Methods). Direct injection detected 3.47% and 0.91% of unlabeled hexanoic and dodecanoic acids respectively, together with mono¹⁸O (m/z +2.004) in 41.85% and 15.89% as well as di¹⁸O (m/z +4.008) isotopologues, confirming more than 95% of incorporation of one to two ¹⁸O atoms.

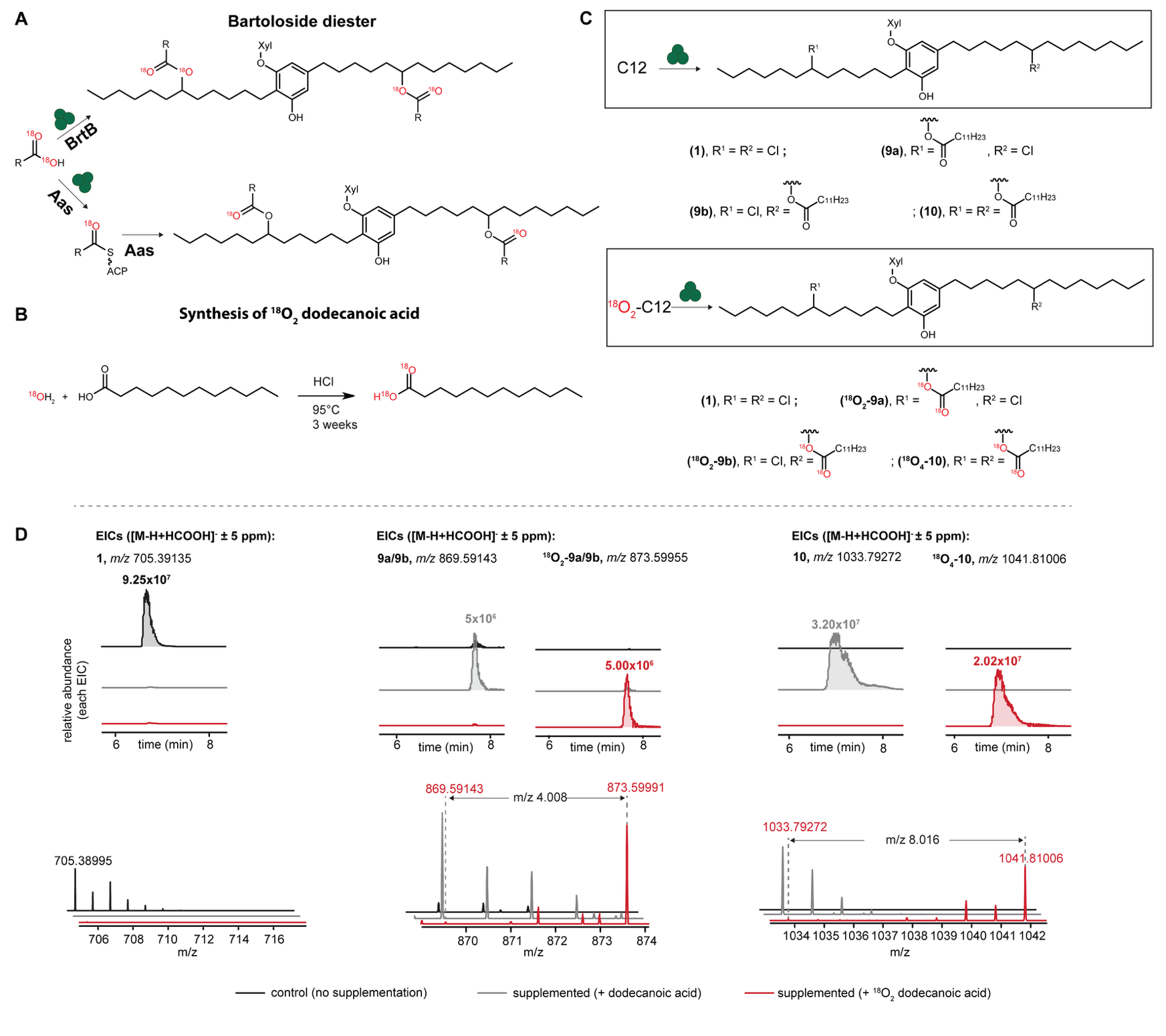

**Figure supplement 2.** **Supplementation of *Synechocystis salina* LEGE 06099 (S06099) with ^18^O_2_-** **dodecanoic acid (C12) leads to the direct esterification of bartolosides into bartoloside fatty acid esters (B-FAs). (A)** Representation of the two different possible pathways taken by the supplemented ^18^O_2_-C12 fatty acid (FA): direct incorporation by BrtB would lead to the incorporation of two (monoester) or four (diester) ^18^O atoms; while prior activation of the FA by the acyl-acyl carrier protein synthetase (Aas) would lead to the incorporation of one (monoester) or two (diester) ^18^O atmos. **(B)** Schematic of the synthesis of ^18^O_2_ dodecanoic acid (C12) from H_2_^18^O and C12. **(C)** Structures of bartoloside A (**1**), mono- and didodecanoate esters (**9a**/**9b**, **10**) and their ^18^O-isotopologues generated upon supplementation of S06099 with ^18^O_2_-C12. **(D)** LC-HRESIMS-based detection of **1**, **9**, **10**, and ^18^O_2 or 4_-labeled **9** and **10** in ^18^O_2_-C12 supplemented cultures of S06099. Values next to each peak in extracted ion-chromatograms (EICs) correspond to peak height (ion counts). ACP, acyl–acyl carrier protein; Xyl, xylose.

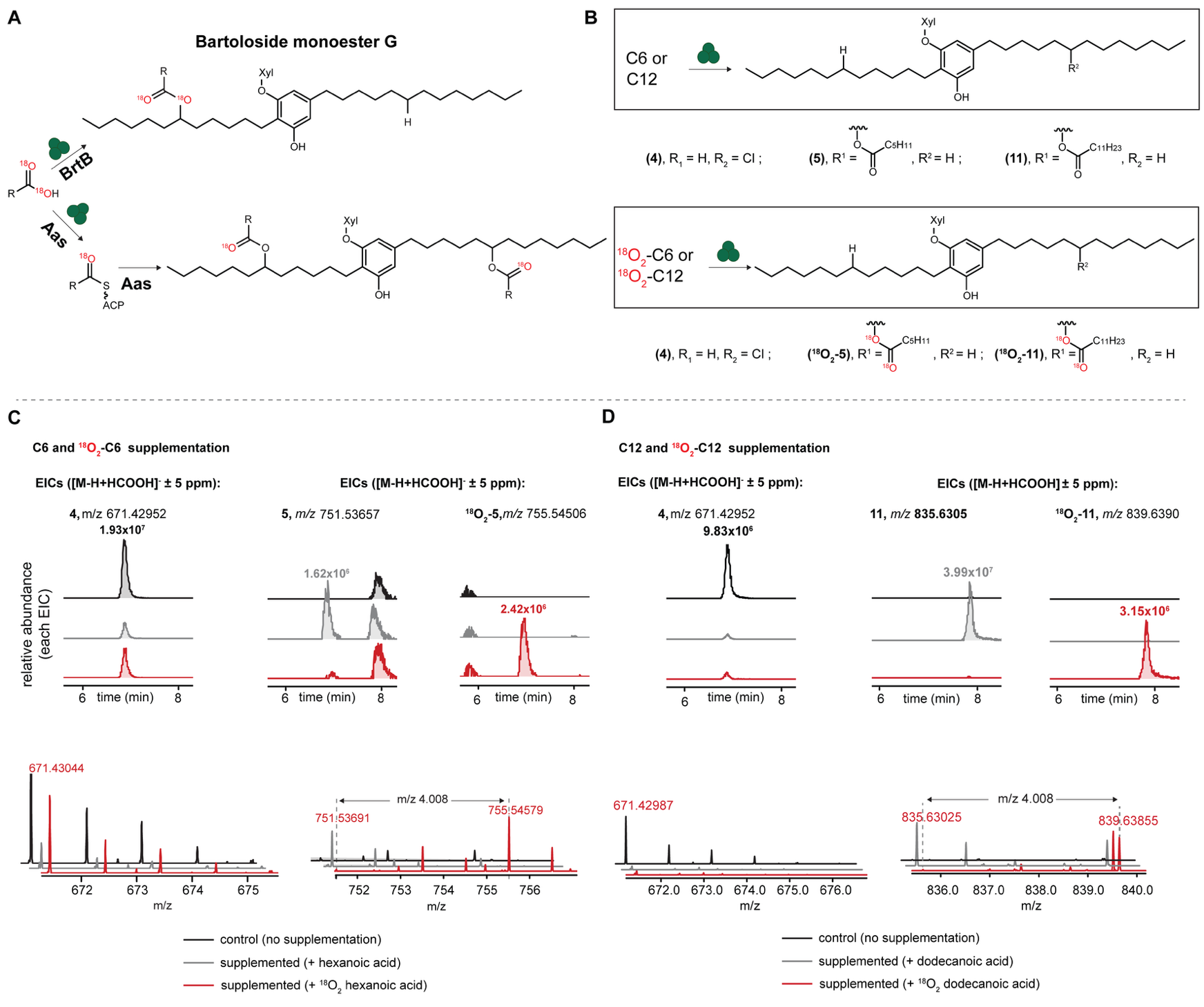

**Figure supplement 3. Supplementation of *Synechocystis salina* LEGE 06099 (S06099) with ^18^O_2_-hexanoic (C6) or with ^18^O_2_-dodecanoic (C12) acids leads to the direct esterification of bartolosides G into bartoloside fatty acid esters (B-FAs) G. (A)** Representation of the two different possible pathways taken by the supplemented ^18^O_2_-C12 fatty acid (FA): direct incorporation by BrtB would lead to the incorporation of two (monoester) ^18^O atoms, while prior activation of the FA by the acyl-acyl carrier protein synthetase (Aas) would lead to the incorporation of one (monoester) ^18^O atom. **(B)** Structures of bartoloside G (**4**), monohexanoate (**5**) monododecanoate (**11**) esters, and their ^18^O-isotopologues generated upon supplementation of S06099 with ^18^O_2_-C6 or ^18^O_2_-C12. **(C)** LC-HRESIMS-based detection of **4**, **5** and ^18^O_2_ labeled **5** in ^18^O_2_-C6 supplemented cultures of S06099. **(D)** LC-HRESIMS-based detection of **4**, **11** and ^18^O_2_ labeled **11** in ^18^O_2_-C12 supplemented cultures of S06099. Values next to each peak in extracted ion-chromatograms (EICs) correspond to peak height (ion counts); ACP, acyl–acyl carrier protein; Xyl, xylose.

**Source data 1 (excel). Extracted** **LC-MS peak area (area under the curve) of bartoloside-related metabolites after fatty acid (FA) supplementation.** Samples 1–3, 5–7 and 9–11 were supplemented with 0.01, 0.05 and 0.5 mM of deuterated FA (*d*_15_C8 m/z + 15,09415 for one labeled chain and + 30.1883 for two labeled chain ; *d*_23_C12 m/z + 23.14426 for one labeled chain and m/z + 46.288 for two labeled chain) respectively, and samples 4, 8 and 12 received DMSO only. T1 samples were collected 24 h after supplementation and T2 samples were collected 7 days after supplementation. Di esters labeled species were detected with either one labeled FA chain (“*d*15, *d*23”) or two labeled FA chains (“*d*30, *d*46”).

SourceData_1_C12_C8_supplementation_bartolosides_related_metabolites;

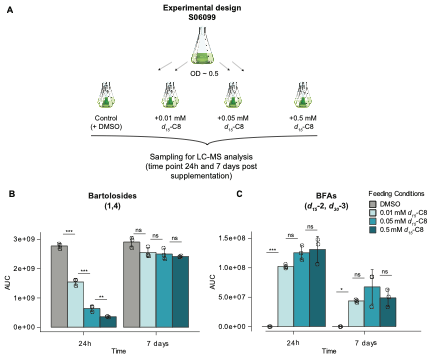

**Figure supplement 4.** ***Synechocystis salina* LEGE 06099 (S06099) bartoloside levels decrease with increasing supplementation of *d*_15_–octanoic acid (C8) while bartoloside fatty acid esters (B-FA) levels plateau. (A)** Design of labeled fatty acid (FA) dose-response experiment. Mid-exponential cultures (OD ~0.5) of S06099 were supplemented with 0.01, 0.05 or 0.5 mM *d*_15_–C8 (or DMSO as a control) and harvested after 24 h and 7 days for LC-HRESIMS analysis. **(B)** Sum of area under the curves (AUCs) for **1** and **4**. **(C)** Sum of AUCs for *d*-labeled octanoate esters (BFAs; *d*_15_-**2**, *d*_30_-**3**). Bars show mean ± s.d. (n = 3 biological replicates); asterisks indicate significant differences between conditions (ns, not significant; * P < 0.05; ** P < 0.01; *** P < 0.001). All AUC values correspond to extracted ion chromatograms of the [M–H+HCOOH]^⁻^ formate adduct of the indicated species. S06099, *Synechocystis salina* LEGE 06099, AUC, area under the curve, DMSO, dimethyl sulfoxide; LC-MS, liquid chromatography-mass spectrometry.

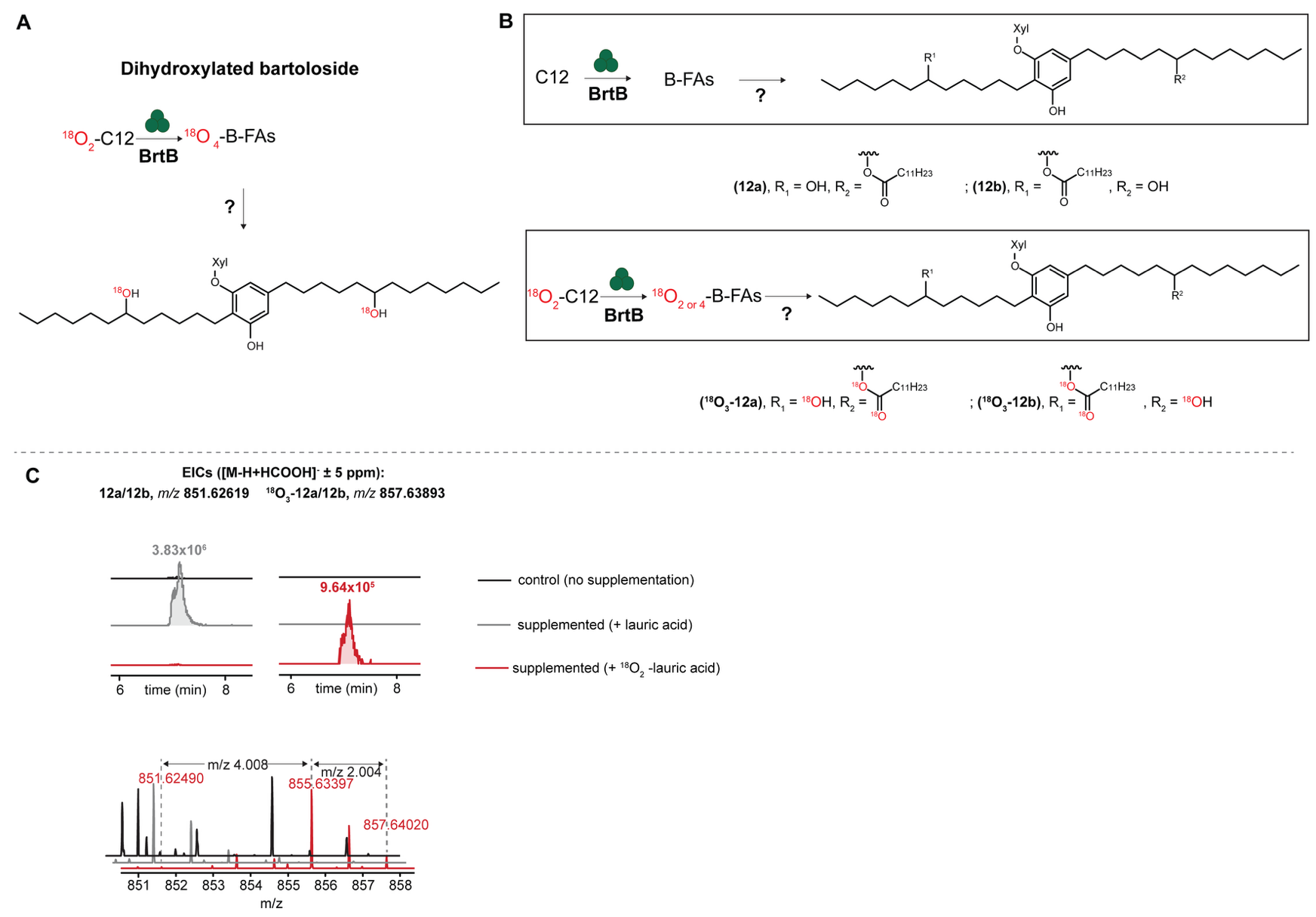

**Figure supplement 5.** **Bartoloside fatty acid esters (B-FAs) are hydrolyzed to hydroxybartolosides (B-OHs). (A)** Representation of proposed events following ^18^O_2_-dodecanoic acid (C12) supplementation: direct esterification by BrtB leading to the incorporation of two or four ^18^O atoms (two per each ester bond), followed by subsequent hydrolysis of the ester bonds to generate dihydroxylated bartolosides containing one or two ^18^O labeled B-OH. **(B)** Structures of B-OHs (**12**) and their ^18^O-isotopologues generated upon supplementation of *Synechocystis salina* LEGE 06099 (S06099) with ^18^O_2_-C12. **(C)** LC-HRESIMS-based detection of **12** and ^18^O_3_ labeled **12** in ^18^O_2_-C12 supplemented cultures of S06099. Values next to each peak in the extracted ion-chromatograms correspond to peak height (ion counts). Xyl, xylose.

**Source data 2 (excel). Extracted** **LC-MS peak areas (area under the curve) of phosphatidylglycerols (PGs) and sulfoquinovosyldiacylglycerols (SQDGs) after fatty acid supplementation.** Samples 1–3, 5–7 and 9–11 were supplemented with 0.01, 0.05 and 0.5 mM of deuterated FA (*d*_15_C8 m/z + 15,09415 for one labeled chain and + 30.1883 for two labeled chain ; *d*_23_C12 m/z + 23.14426 for one labeled chain and m/z + 46.288 for two labeled chain) respectively, and samples 4, 8 and 12 received DMSO only. T1 samples were collected 24 h after supplementation and T2 samples were collected 7 days after supplementation. PG- and SQDG-associated labeled species were search with either one labeled FA chain (“*d*15, *d*23”) or two labeled FA chains (“*d*30, *d*46”) and 0 to 4 unsaturations.

SourceData_2_C12_C8_ supplementation_PGs_SQDGs

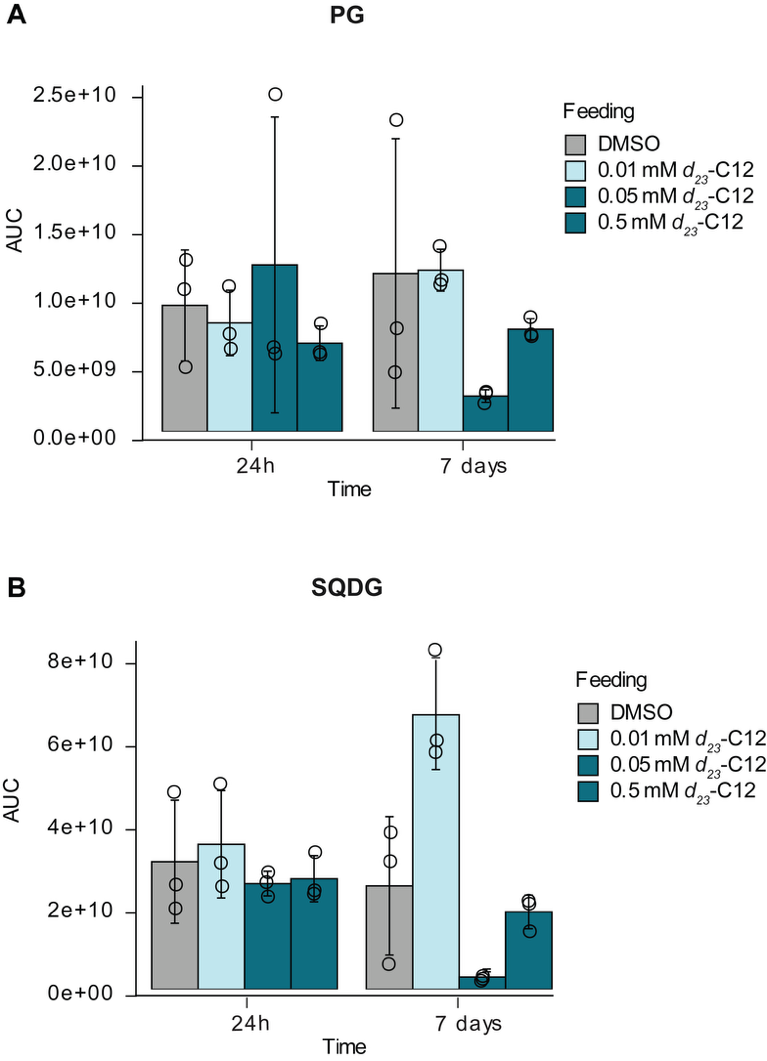

**Figure supplement 6.** **Fatty acid supplementation does not change non-labeled glycolipid levels.** Extracted LC-MS peak areas (AUC) of non-labeled phosphatidylglycerols (PG; A) and sulfoquinovosyldiacylglycerols (SQDG; B) in *Synechocystis salina* LEGE 06099 after supplementation with *d₂₃*–C12 (0.01, 0.05, or 0.5 mM; in DMSO) or DMSO alone. Samples were collected 24 h and 7 days after supplementation. Bars show mean ± s.d., with individual biological replicates overlaid (n = 3). AUC values correspond to extracted ion chromatograms of the [M–H+HCOOH]^⁻^ formate adducts. AUC, area under the curve; DMSO, dimethyl sulfoxide.

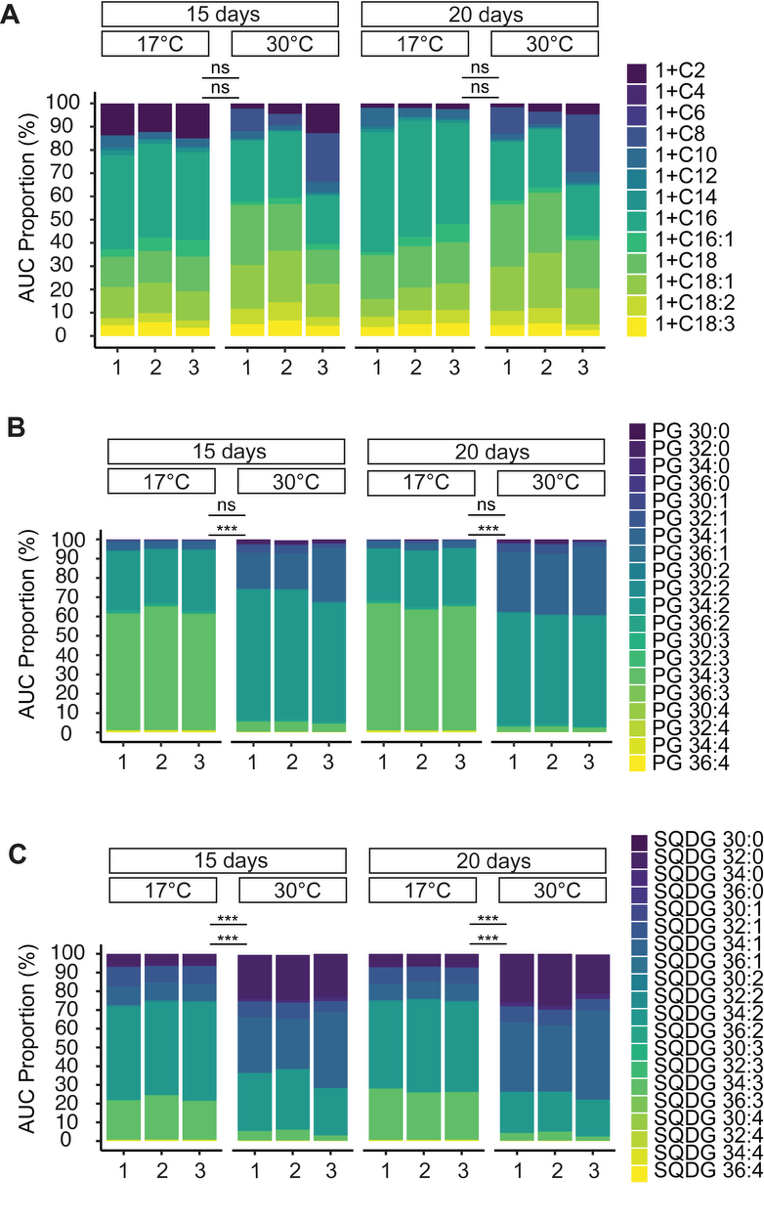

**Figure supplement 7.** **Bartoloside fatty acid ester (B-FA) acyl composition is stable, while anionic glycerolipids remodel with temperature.** Sum of area under the curves (AUCs) for B-FAs (A), **phosphatidylglycerols (**PGs) (B) and **sulfoquinovosyldiacylglycerols (SQDGs). Anionic glycerolipids** here refers to **PGs** and **SQDGs** quantified by negative-mode LC-MS; other lipid classes were not assayed. Sampling days were chosen to include one point near **OD ~0.5** at each temperature, to compare **matched physiological states**. “1”, “2”, “3” refers to the different triplicates in each condition. For each triplicate, the AUC per compound was extracted and then transformed into proportion. Asterisks indicate significant temperature effects (17°C vs 30°C) on replicates-level indices (B-FA/PG/SQDG long-chain content (top) and unsaturation content (bottom); Welch’s t-test with Benjamini–Hochberg correction, p < 0.05). “ns” marks non-significant contrasts and “***” marks p < 0.001.

**Source data 3. Extracted** **LC-MS peak areas (area under the curve) of bartolosides fatty acid esters, PGs and SQDGs observed at 17°C and 30°C in S06099.**

Source_Data_3_temperature_AUC.

**Source data 4:** **Optical Density (OD) of *Synechocystis salina* LEGE 06099 (S06099) cultures used for the transcriptomics and metabolomics assay and extracted** **LC-MS peak areas (AUC) of bartoloside related metabolites.**

Source_data_4_OD_metabolomics

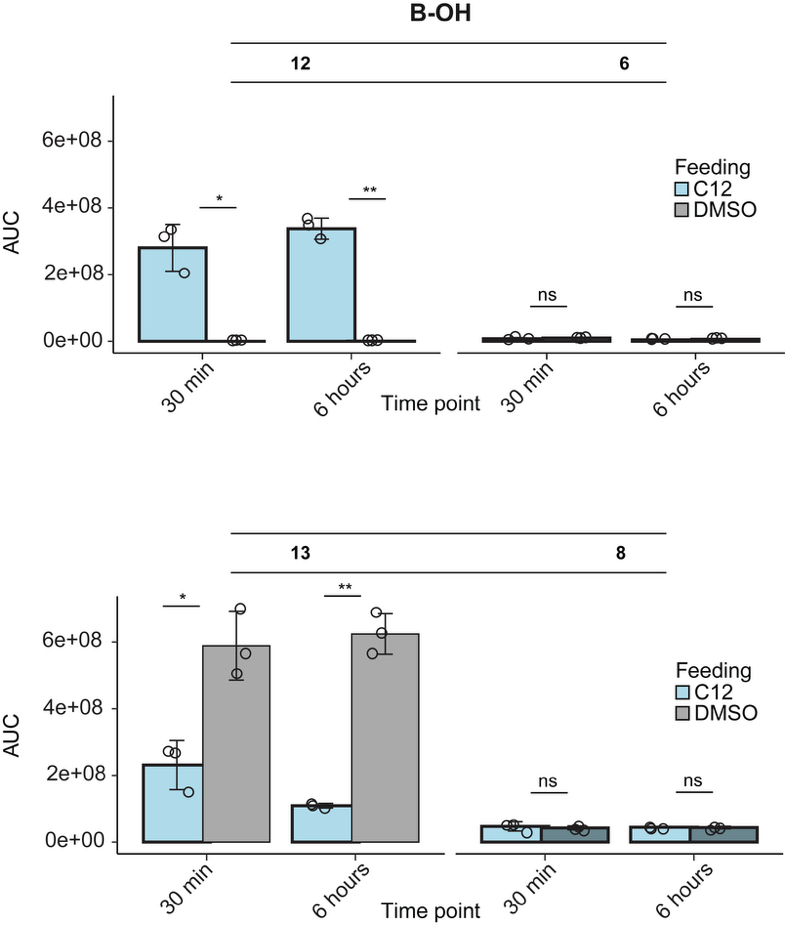

**Figure supplement 8.** **Hydroxy-bartoloside (B-OH) profiles grouped by type.** Total LC-MS peak area (AUC) of each B-OH type, obtained by summing separately the AUC of all detected species containing that hydroxyl group (**13a/b**, **12a/b**, **6**, **8**), at 30 min and 6 h in dodecanoic-acid-treated vs control cultures; bars show mean ± s.d. for biological replicates and asterisks indicate statistical significance (* P < 0.05; ** P < 0.01; ns, not significant). B-OH, hydroxy bartoloside; AUC values correspond to extracted ion chromatograms of the [M–H+HCOOH]^⁻^ formate adducts. AUC, area under the curve; DMSO, dimethyl sulfoxide.

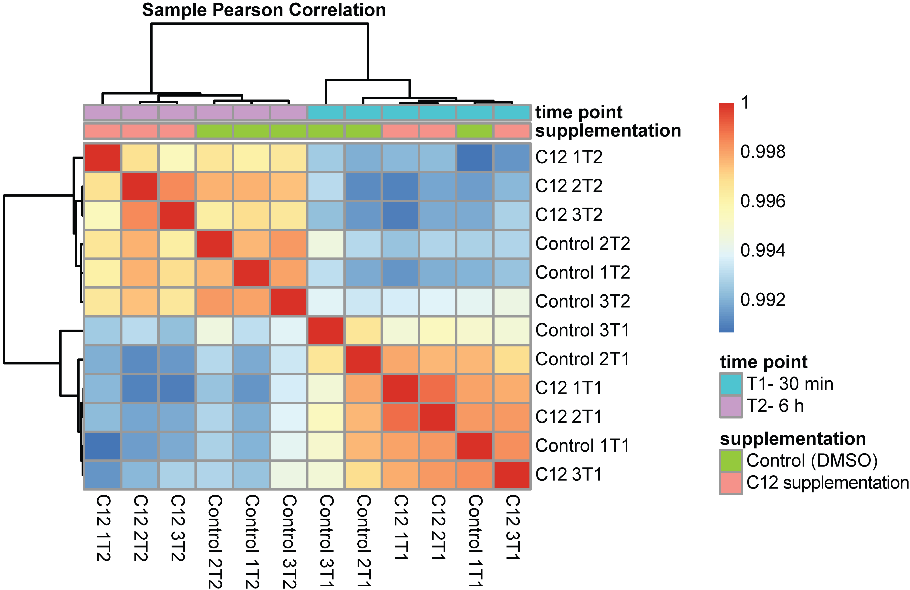

**Figure supplement 9.** **Sample-wise Pearson correlation of RNA-seq libraries.** Heatmap showing pairwise Pearson correlation coefficients between all RNA-seq samples, calculated from normalized gene expression values. Samples cluster primarily by timepoint (T1, 30 min; T2, 6 h), with additional separation by treatment at 6 h, revealing near-identical expression profiles across all libraries (r > 0.99), and no sample deviated from this pattern. Color scale indicates the Pearson correlation coefficient (r); DMSO, dimethyl sulfoxide.

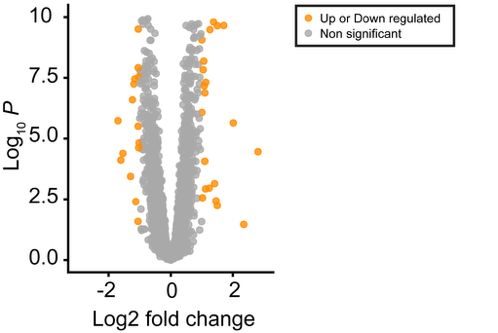

**Figure supplement 10. Time-dependent gene expression in *Synechocystis salina* LEGE 06099 (S06099).** Volcano plot of differential expression for 6 h versus 30 min samples in S06099, testing the effect of time. Points show log_2_ fold change (6 h/30 min) versus –log_10_ adjusted p value (FDR); genes meeting |log_2_FC| ≥ 1 and FDR ≤ 0.05 are highlighted as significantly up- or downregulated.

**Table supplement 1 (excel).** **Differentially expressed genes between 30 min and 6 h regardless of treatment** (Table_supplement_1_DE genes_time).

**Text supplement 1.** **BrtB amino acid sequence and annotated features.**

MANPFDVALTNLNSLYSGIPDVPVLQTYNSLNPYDISLTNLNNLSNFVIEPAVLPTAAGTYGPQISEIAFALGGGIITGALADLLSTELGIEIENEALITKFLSFWTPGSDYAFGQDYVEFIQGREGFDTIIGYDPGFNSALAPVQIDFVLGGPRQDFNFFAPLDEPARYVLGDWRTPYYVDNDIQSEGLNEFMYVSLFTLNPDEFSPDNAENHRIQLYGSLDNYYFVPTTVVIDGSALDPSLPSIPVPGTEIYYKTGQVFADLIAIVPFQDPNDPNLISAWIDFEGYLPPEQADLSGQTANPLLSGSPLLQLGYEGIDMGTATALAPDGDIYVAGSTSSLLGPFPRGGSDNFIARYNDDGSLEWIVQFGSTEFDIITDIVSDSAGNVYATGWTRGNVETGFGLAPGVQDNWVAKFDANGSQLWVEQFTIDGYLDRSMGIALDEVNGRIHLTGHSNTDGSVADTNDPAIVPNGNVVTWIAGFDANTGAEDYRNVFEVAQTSNSRGRIDEGFGIDVDDAGNVFSTGWAGENAYDVYLVKSDTDGAVLWTKSFGTGSNATQSYAWDVASDGTNAYILGWTQGELNTFRDQIPPTDEFAGNATAPLTPQNAYQGGTFDAFVAAYDTDGTELWTWHVGGTGDDGTFFGKIVAAGDFVYATGYTDGFIGTLGGSNAGDYDAWIGKFDKLTGQTAWIQQIGSTKLDYATGISVDGDDIFVTGFTEGSLGGLNGGASDAWVAKLNQDGALEAFNVSAPIAPPAPFAPPAPLTPPAPPAPDLSSLQSLGSTPQPTYPALGDSYGTNPFGTQPTQPTYDLSSMYQNAYYGGFGGYSGYGY

BrtB from S06099 is shown with three annotated regions: a glycine/aspartate-rich predicted calcium-binding motif consistent with RTX-like repeats (underlined grey), a proline-rich region proposed to disrupt β-sheet formation (blue), and the C-terminal glycine/serine-rich linker (green). BrtB is 88.6 kDa with a predicted pI of 3.73, contains ~12% glycine, and lacks cysteine residues.

**Source data 5**. Total exoproteome of *Synechocystis salina* LEGE06099 (S06099) without and with C16 supplementation.

**Source data 6-9 (excel).** **Proteome tables for gel band pieces F1-F4** (SourceData3_Fig4_F1; SourceData4_Fig4_F2 ; SourceData5_Fig4_F3 ; SourceData6_Fig4_F4).

**Table supplement 2 (excel).** **Top 10 non-contaminant proteins per gel band piece (LC-MS/MS).** For each excised SDS–polyacrylamide gel band piece (F1–F4), the table lists the **ten most abundant protein entries** after excluding proteins flagged as **Contaminant** in the proteome export (Source data 3-6). Abundances correspond to Source data 3-6 **Abundances (Normalized)** for the indicated sample. “Rank (non-contam)” is the rank within that gel piece among non-contaminant proteins. “Coverage (%)” and “Peptides” report the sequence coverage and the number of identified peptides assigned to each protein, as reported in the Source data 3-6.

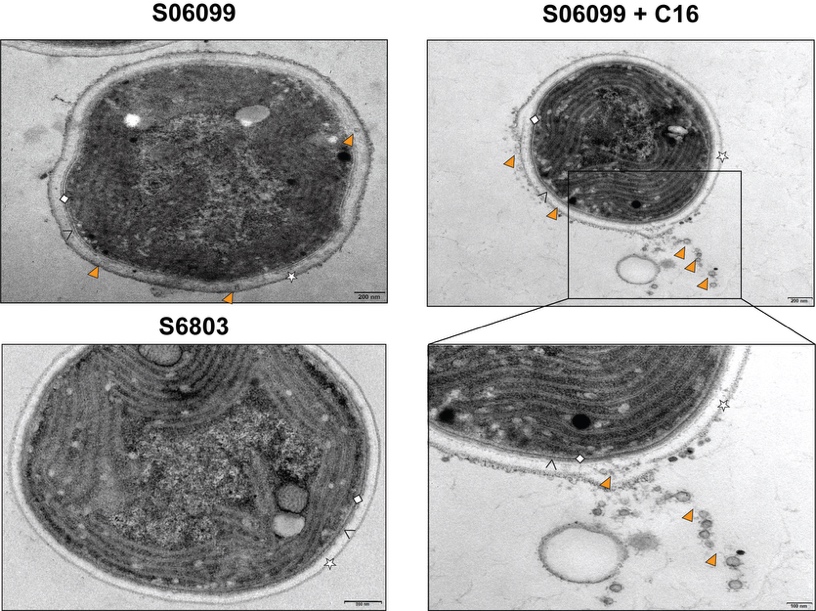

**Figure supplement 11.** **Electron microscopy of *Synechocystis salina* LEGE 06099 with and without FA supplementation.** Transmission electron micrographs of S06099 cells grown without (S06099) and with (S06099 + C16) C16 fatty acid supplementation, and of *Synechocystis* sp. PCC 6803 (S6803) grown without fatty acid supplementation, showing the S-layer (star), outer membrane (chevron) and the peptidoglycan (diamond). Vesicle-like protrusions (arrowheads) at the cell surface can be observed across conditions in S06099. Scale bars, 200 nm (top panels) and 100 nm (bottom-right zoom).
